# Network spreading and local biological vulnerability in amyotrophic lateral sclerosis

**DOI:** 10.1101/2024.04.11.588760

**Authors:** Asa Farahani, Justine Y. Hansen, Vincent Bazinet, Golia Shafiei, D. Louis Collins, Mahsa Dadar, Sanjay Kalra, Alain Dagher, Bratislav Misic

## Abstract

Amyotrophic Lateral Sclerosis (ALS) is a progressive neurodegenerative disease that predominantly targets the motor system. Spread of pathology is thought to be driven by both local vulnerability and network architecture. Namely, molecular and cellular features may confer vulnerability to specific neuronal populations, while synaptic contacts may also increase exposure to pathology in connected neuronal populations. However, these principles are typically studied in isolation and it remains unknown how local vulnerability and network spreading interact to shape cortical atrophy. Here we investigate how network structure and local biological features jointly shape the spatial patterning of atrophy in ALS. We analyze the Canadian ALS Neuroimaging Consortium (CALSNIC) dataset and estimate cortical atrophy using deformation-based morphometry (DBM). We find that structural connectivity closely aligns with the course of atrophy. Atrophy is also more likely to occur in regions that share similar transcriptomic, neurotransmitter receptor and metabolic profiles. We identify disease epicenters in motor cortex. Epicenter probability maps show transcriptomic enrichment for biological pathways involved in mitochondrial function as well as support cells, including endothelial cells and pericytes. Finally, individual differences in epicenter location correspond to individual differences in clinical and cognitive symptoms, and differentiate patient subtypes.

## INTRODUCTION

Amyotrophic Lateral Sclerosis (ALS) is a terminal neurodegenerative disease associated with progressive impairment of motor functions [30]. Expected survival of most patients diagnosed with ALS is 2–5 years after onset [4, 86]. The disease shows significant heterogeneity across individuals, and patients can be classified into subtypes based on multiple features, including the familial or sporadic occurrence of the disease, age of disease onset, symmetry of motor neuron involvement, and initial symptom location [129]. Most ALS patients experience a progression of symptom severity, which reflects ALS pathology spread in the central nervous system.

Most modern accounts of ALS pathology revolve around two non-exclusive notions. The first is network spreading: that the spread of pathology, likely in the form of pathogenic misfolded proteins, occurs via synaptic contacts [20, 21, 89, 94, 97, 98, 112, 142, 149]. Here initial infiltration via the corticospinal tract introduces pathogenic proteins in primary motor cortex, leading to progressive cell death and atrophy. The second notion is that of local vulnerability: that molecular and morphological features of specific cells predispose them to the disease [108, 121]. In addition, the involvement of support cells has been noted in ALS, including astrocytes and pericytes, implicating energy homeostatic and vascular mechanisms [11, 81, 107, 109]. Importantly, the two perspectives may both be true; namely, pathology may spread via synaptic contacts, but the spread may be amplified in vulnerable neuronal populations, effectively guiding the network spread of pathology.

What are the principles that shape the spatial patterning of atrophy in ALS? Here we address this question using the Canadian ALS Neuroimaging Consortium (CAL-SNIC) dataset (http://calsnic.org) [66]. We first establish that cortical atrophy reflects white matter architecture. We also assess the extent to which the spread of atrophy between brain regions depends on their molecular and cellular similarity, including transcriptomic similarity, neurotransmitter receptor similarity, and metabolic similarity. We then use methods from epidemiology to identify network epicenters of the disease process in the cortex. We show that cortical epicenters of pathology co-localize with markers of metabolic and mitochondrial function. Finally, we show that individual differences in epicenter location can distinguish subtypes of patients (bulbar-versus spinal-onset) and correlate with clinical and cognitive function.

## RESULTS

Data were derived from the CALSNIC repository [66], and comprised *N* = 192 patients and *N* = 175 healthy age- and sex-matched control participants. Atrophy was estimated using deformation-based morphometry (DBM), a morphometric technique that has previously been shown to be sensitive to tissue loss in both deep and superficial structures [9], and across multiple neurodegenerative syndromes [17, 36, 119, 132, 134]. For details on data acquisition or preprocessing, see *Methods*.

### Spatial distribution of atrophy

We initially identify differences between the ALS patients and healthy controls. Fig. 1a shows the group-average atrophy map. Throughout the manuscript, we use sign-inverted *w*-scores such that greater values correspond to greater atrophy: a *w*-score is a morphometric measure of atrophy that is corrected for differences in age, sex and imaging site. Consistent with previous reports, the map highlights pronounced atrophy throughout the brain, including both grey matter (cortex and subcortex) and white matter [36].

**Figure 1.**
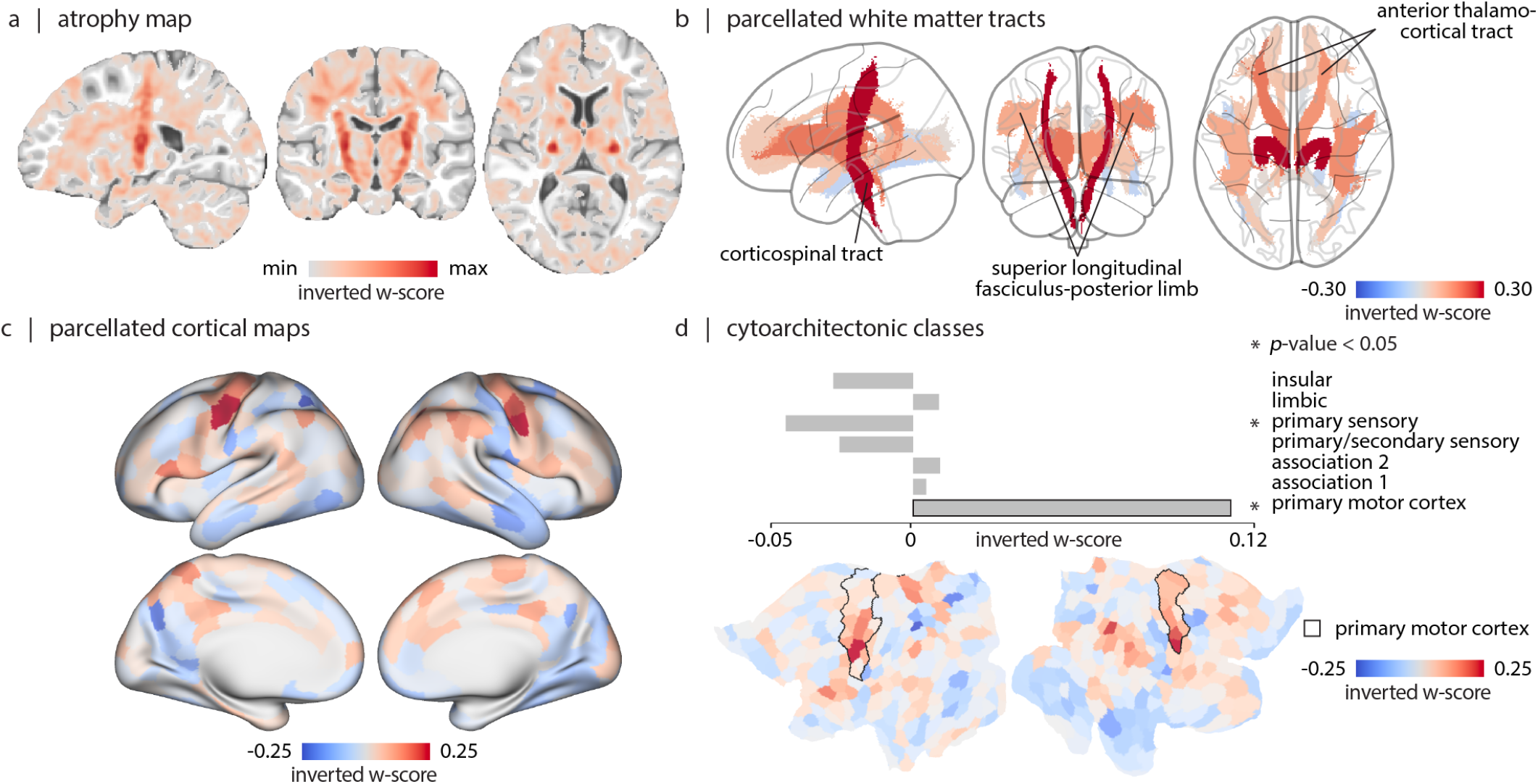
ALS-related atrophy. (a) Sagittal, coronal, and axial view of mean atrophy across individuals with ALS. Atrophy is estimated as the group-average *w*-score across individuals with ALS. The map is thresholded to exclude voxels with atrophy values smaller than 0. Data are displayed on a T1-weighted MNI template (MNI152-NonLinear2009cSym - 1 × 1 × 1*mm*; *P* = 118, *C* = 117, *A* = 87). (b) Sagittal, coronal, and axial view of white matter tract parcels colored by mean atrophy. White matter tracts are defined based on the JHU white matter tractography atlas (thresholded at 25% probability) [60, 146]. Significant atrophy is observed in bilateral corticospinal tracts (FDR corrected; left: *p* = 8.83 × 10^−16^, *t*-statistic= 9.62; right: *p* = 4.13 × 10^−16^, *t*-statistic= 9.84), anterior thalamocortical tracts (FDR corrected; left: *p* = 6.53 × 10^−8^, *t*-statistic= 6.44; right: *p* = 2.04 × 10^−5^, *t*-statistic= 5.26) and superior longitudinal fasciculus-posterior limb tracts (FDR corrected; left: *p* = 7.39 × 10^−4^, *t*-statistic= 4.34; right: *p* = 1.68 × 10^−4^, *t*-statistic= 4.75). (c) Cortical atrophy map parcellated based on Schaefer-400 parcellation [111]; maps are displayed on fs-LR inflated cortical surfaces. (d) Mean atrophy is calculated within each of the cytoarchitectonic classes defined by Von Economo [39, 113, 145]. Statistical significance is estimated using a spatial autocorrelation-preserving spin test (1, 000 repetitions). Black borders shown on the fs-LR flat cortical surface correspond to the primary motor cortex borders defined by Von Economo cytoarchitectonic parcellation [39, 113, 145].

Given that most cortical atrophy is concentrated in primary motor cortex, we implement an initial sanity check: whether atrophy can correspondingly be observed in the corticospinal tract. We segment the voxel-wise atrophy map using the Johns Hopkins University (JHU) white matter tractography atlas [60, 146] (Fig.1b). Analysis of white-matter tracts reveal significant involvement of bilateral corticospinal tract, bilateral anterior thalamic radiation, and bilateral superior longitudinal fasciculus bundles (*p <* 0.05, False Discovery Rate (FDR) corrected; Fig. 1b). These projections have previously been associated with ALS and suggest that cortical atrophy is related to white matter atrophy [3, 18, 37, 90, 104]. In the subsequent analyses we directly assess the relationship between local grey matter atrophy and network connectivity.

To cross-reference the atrophy map with network connectivity data, we apply a high resolution functional parcellation, subdividing the atrophy map into 400 parcels according to the Schaefer atlas [111] (Fig. 1c). In the cortex, the most pronounced atrophy is observed near the pre-central gyrus, a hub for motor function. Atrophy also extends into the temporal and frontal cortices (Fig. 1c). To test whether the disease selectively targets primary motor cortical neurons, we compute mean atrophy in each of the canonical cytoarchitectural classes according to the histological Von Economo atlas [39, 113, 145] (Fig. S1). We observe significant enrichment of atrophy in the primary motor cytoarchitectonic class (FDR corrected, *p*_spin_ = 6.99×10^−3^; Fig 1d). Collectively, these results suggest that the present morphometric approach is sensitive to the pathophysiology of ALS.

### Structural connectivity shapes cortical atrophy

We next assess the extent to which the spatial patterning of atrophy is related to structural connectivity. We compute the correlation between a node’s atrophy value and the mean atrophy of its structurally connected neighbours, weighted by streamline density estimated using diffusion MRI (Fig. 2a). To ensure that connectivity estimates reflect the healthy connectome prior to disease onset and deafferentation, we estimated structural connectivity in a sample of *N* = 326 healthy young adults from the Human Connectome Project (S900-HCP; [136]). Fig. 2a shows a positive correlation between the two (Pearson correlation coefficient; *r* = 0.46), suggesting that pathology in a brain region is correlated with greater exposure to pathology in anatomically connected regions, an effect that has been demonstrated in other neurodegenerative syndromes [56, 117, 119, 120, 153].

**Figure 2.**
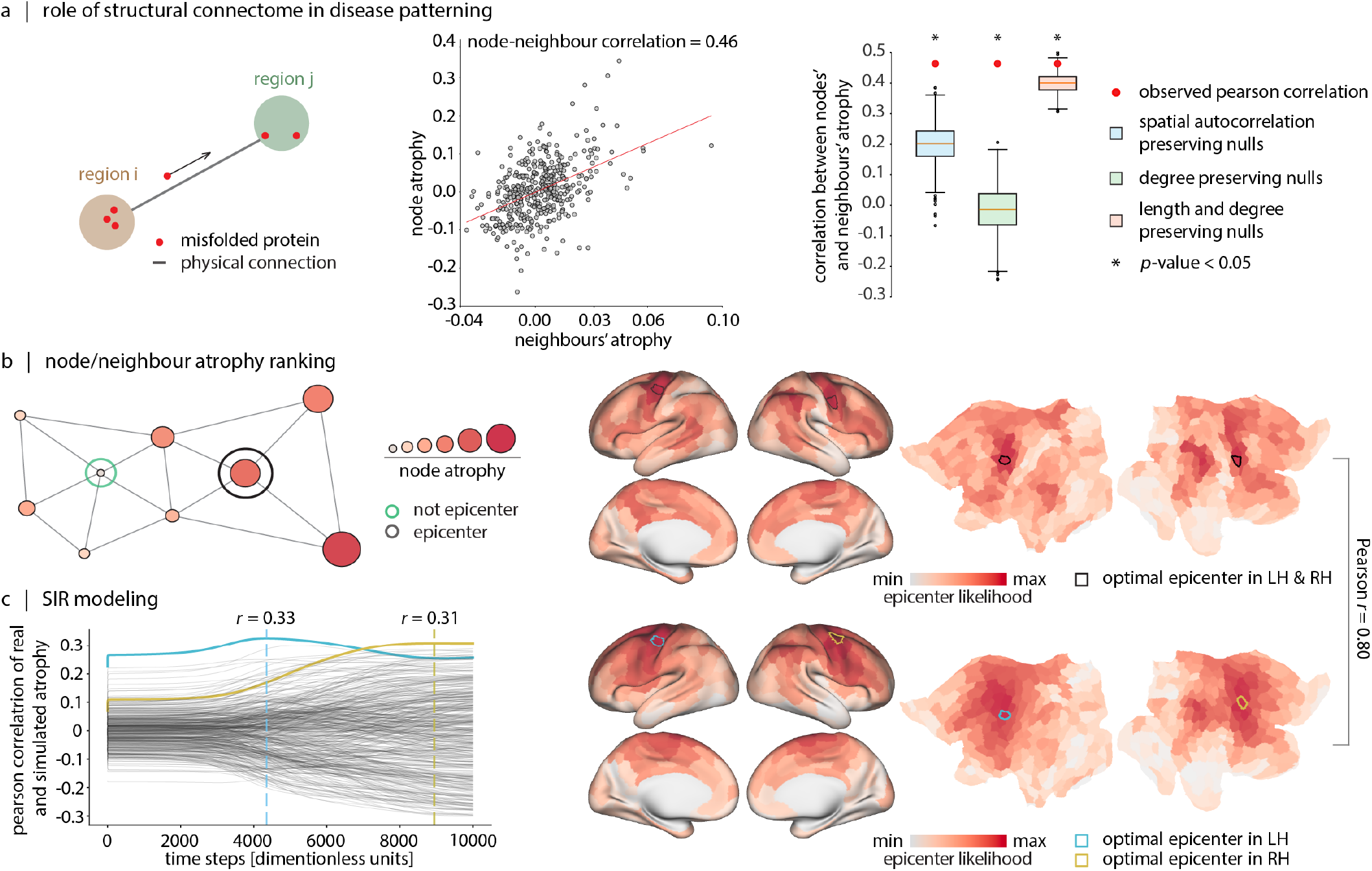
Structural connectivity shapes ALS-related atrophy. (a) Left: We test the hypothesis that local regional atrophy is related to atrophy in its anatomically connected neighbours. The scatter plot shows the atrophy of a node (y-axis) and the mean atrophy of that node’s structurally connected neighbours (Pearson correlation coefficient; *r* = 0.46). Grey circles indicate brain regions. Right: The observed correlation coefficients between node and neighbour atrophy (red circles) are shown with respect to three null models: (1) spatial autocorrelation preserving spin tests (*p* = 1.99 × 10^−3^, blue box plot), (2) degree-preserving rewired networks (*p* = 9.99 × 10^−4^, green box plot), and (3) degree- and edge length-preserving rewired networks (*p* = 0.045, red box plot). Asterisks indicate statistical significance with respect to each null model. (b) In the atrophy ranking method, epicenters are defined as nodes that are both highly atrophied and structurally connected to other highly atrophied nodes. To estimate the epicenter likelihood of a node, nodes are first ranked according to their atrophy and then ranked according to their neighbours’ atrophy. The epicenter likelihood ranking of each node is defined as its mean ranking in the two lists. Visualizations of the epicenter maps, including the top-ranked nodes distinguished by black borders, are presented on both inflated and flat cortical fs-LR surface representations. For completeness, we additionally plot the mean and standard deviation of epicenter maps across individual ALS patients in Fig. S2. (c) Agent-based SIR model locates similar cortical epicenters. The model only considers the structural connectome as the underlying network for pathology spread. The spreading process is initiated in every brain region and the correlation between the simulated and empirical patterns of atrophy is computed at each simulation time point. The two largest correlations are obtained by seeding regions within the motor cortex (indicated by a blue border on the left cortical surface (LH) and by a yellow border on the right cortical surface (RH)). The epicenter likelihood maps obtained by the SIR modeling approach are shown on both inflated and flat fs-LR cortical surfaces. The epicenter likelihood maps obtained via both the ranking method and the SIR modeling lead to cortical patterns which are correlated with each other (Pearson correlation coefficient; *r* = 0.80, *p*spin = 9.99 × 10^−4^, *n*spin = 1, 000).

To demonstrate that atrophy patterns are mainly driven by network topology and not by spatial proximity among regions or the spatial autocorrelation of the atrophy patterns, we apply three null models [138]. The first null model is a spatial autocorrelation-preserving randomization that tests whether the correlation between node and neighbour atrophy is passively due to spatial autocorrelation in the atrophy map (“spin test”; [5, 84]. This model generates a null distribution for node-neighbour correlation values by projecting the atrophy map to a sphere, applying random angular rotations, bringing the rotated map back to the cortical surface, and re-calculating the node-neighbour atrophy correlations using the values from the rotated atrophy maps. We observe significantly greater correlations for the empirical map compared to the rotated maps (*p*_spin_ = 1.99 × 10^−3^), suggesting that the effect cannot be attributed to spatial autocorrelation. Moreover, the correlation between node and neighbour atrophy is also significantly greater when using the empirical structural brain network compared to rewired null structural networks that randomize network topology, including both degree-preserving and degree- and edge length-preserving nulls (*p* = 9.99 × 10^−4^, *p* = 0.045; respectively) [14]. Collectively, these null models show that the correlation between node and neighbour atrophy is specifically due to network topology.

### Epicenters of cortical atrophy

Given that the structural connectome shapes the spread of ALS-related pathology, we next sought to identify the putative cortical epicenter of the pathology. We apply two methods – one empirical and one computational – to back-reconstruct the spreading trajectory and infer the most likely cortical location of the epicenter: (1) a network-based node ranking method, and (2) a susceptible-infected-removed (SIR) dynamical model. Both methods have previously been applied to understand the course of multiple neurological diseases [1, 54, 55, 119, 120, 157]. The ranking method identifies an epicenter as a region that is severely impacted by disease-related atrophy, and whose structurally connected regions also exhibit extensive atrophy. In this approach, brain regions are ranked based on both their own atrophy values and their neighbours’ atrophy values in two separate lists. Each cortical node is then assigned a value reflecting the node’s average rank across these two lists. Fig. 2b showcases the final mean ranking values across brain regions; in this map, higher values indicate higher probability of a node being an epicenter. The two most probable epicenter locations are within the right and left pre-central gyrus (primary motor cortex).

In the second approach, we build an SIR model using the structural connectome as the only underlying foundation for the spread of pathogenic agents, and hence atrophy. The model works by simulating the misfolding of normal proteins in the cortex and their transneuronal spread through the structural connections between brain regions. The similarity between real atrophy maps and simulated atrophy maps is computed at each time point (Fig. 2c). Each trajectory in the plot corresponds to the similarity of real and simulated atrophy when a specific parcel is chosen as the epicenter for the spread of misfolded proteins. Brain parcels are ranked based on parcels’ maximal correlation value across all simulation time-points. The top two parcels which can best reproduce the atrophy pattern, are indicated in blue and yellow in Fig.2c, and again are located within the motor cortex. Altogether, both methods yield similar epicenter probability maps (Pearson correlation coefficient; *r* = 0.80, *p*_spin_ = 9.99 × 10^−4^, *n*_spin_ = 1, 000). Both maps suggest high probability of being an epicenter for parcels across the motor and premotor cortices, as well as dorsolateral prefrontal cortex, posterior parietal cortex, and superior temporal gyrus.

### Local biological features guide pathogenic spread

We next investigate the potential contribution of multiple biological features in guiding disease spread. We consider the hypothesis that, while disease spread occurs via axonal projections, spread may be more likely between regions that display or share specific biological features. Specifically, we reconstruct five inter-regional similarity networks that describe the biological similarity of pairs of brain regions. The networks include: (1) gene expression similarity, (2) neurotransmitter receptor similarity, (3) laminar differentiation similarity, (4) metabolic similarity, and (5) hemodynamic similarity (i.e. functional connectivity) (Fig. 3a) [56].

**Figure 3.**
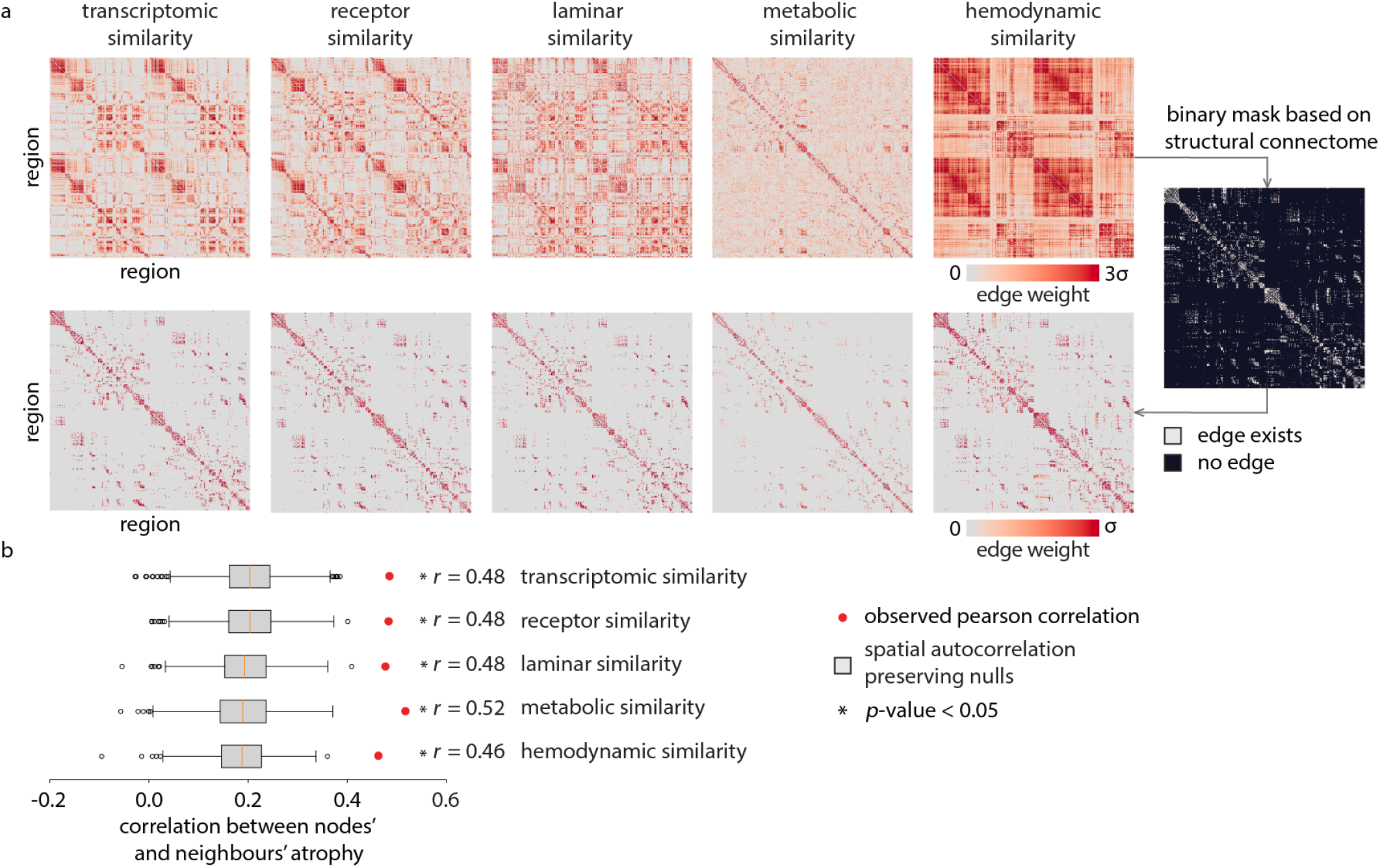
Inter-regional biological similarity shapes atrophy. Inter-regional similarity networks reflect the similarity of brain regions according to multiple biological features. We analyze transcriptomic (gene expression) similarity, receptor similarity, laminar similarity, metabolic similarity, and hemodynamic similarity [56]. (a) The heatmaps in the first row visualize these networks. Negative-valued elements are excluded from all analyses. Similarity matrices are masked based on the structural connectome (using structural connections and weights from each of the inter-regional similarity matrices), resulting in heatmaps shown in the second row. (b) The node-neighbour atrophy correlations are estimated for each masked inter-regional similarity network. The significance of node-neighbour correlations are assessed with respect to spatial autocorrelation preserving spin tests (*n*_spin_ = 10, 000). Red dots depict node-neighbour correlations, and boxplots depict the corresponding spin test-estimated null distributions.

To assess whether regions with similar biological features are also more likely to experience atrophy, we compute node-neighbour similarity as in Fig. 2a. Namely, we define the “exposure” that region *i* has to region *j*’s atrophy as the product between the edge weight (*c*_*ij*_ if *c*_*ij*_ *>* 0) and the extent of atrophy in the neighbouring nodes. Here we use the connection weights derived from each of the five inter-regional similarity networks masked by the structural connectome (using structural connections and weights from each of the inter-regional similarity matrices). If the weights from a network yield a high value for node-neighbour atrophy, this suggests that the biological feature encoded by that network contributes to the spread of pathology [56]. We assess significance with respect to spatial autocorrelation-preserving spin tests and apply FDR correction to the resulting *p*-values.

We find that all networks of inter-regional biological similarity yield significant correlations between node and neighbour atrophy values, suggesting that ALS pathology is more likely to spread between regions with similar biological features (Fig. 3b). Interestingly, metabolic similarity – estimated using dynamic FDG PET – yields the greatest node-neighbour similarity (Pearson correlation coefficient; *r* = 0.52, *p*_spin_ = 1.25 × 10^−3^). This is consistent with numerous reports that ALS is associated with metabolic dysfunction, including abnormal mitochondrial physiology leading to a decreased level of adenosine triphosphate (ATP) and oxidative stress, as well as dysfunction of astrocyte mitochondrial and glutamate transporters leading to increased capture of free glutamate and excitotoxicity [37, 130, 137]. The results show that the addition of biological similarity information results in increased node-neighbour correlation values (gene expression similarity, *r* = 0.48; neurotransmitter receptor similarity, *r* = 0.48; metabolic similarity, *r* = 48; metabolic similarity, *r* = 0.52; hemodynamic similarity, *r* = 0.46) compared to using structural connectivity alone (*r* = 0.46), suggesting that network structure and local biological features jointly contribute to the spatial patterning of atrophy.

For completeness, we repeated the analysis using the unmasked inter-regional similarity matrices (using connectomes shown in the first row of Fig. 3a, and Fig. S3). In this case, we also observe significant node-neighbour atrophy using gene expression similarity (Pearson correlation coefficient; *r* = 0.35, *p*_spin_ = 1.33 × 10^−2^), neurotransmitter receptor similarity (Pearson correlation coefficient; *r* = 0.35, *p*_spin_ = 1.25 × 10^−2^), metabolic similarity (Pearson correlation coefficient; *r* = 0.43, *p*_spin_ = 4.99 × 10^−3^), and hemodynamic similarity (Pearson correlation coefficient; *r* = 0.25, *p*_spin_ = 2.75 × 10^−2^). In the next subsection, we investigate the molecular and cellular features associated with atrophy in greater detail.

### Molecular and cellular signatures of epicenters

Up to now, we find that atrophy patterns reflect network organization and are centered on a compact set of epicenters. We next ask whether the network epicenters of ALS atrophy are enriched for specific molecular pathways and cell types. We cross-reference the ALS epicenter probability map with microarray gene expression from the Allen Human Brain Atlas [57]. We submitted the gene list to Gene Category Enrichment Analysis (GCEA) to isolate Gene Ontology (GO) categories in which the constituent genes are significantly more correlated with atrophy than a population of random atrophy maps with preserved spatial autocorrelation [44] (see *Methods*).

Consistent with the intuition developed in the previous subsection – suggesting involvement of metabolic features in disease spread – the top GO categories are mainly associated with metabolic processes and energetic homeostasis (Fig. 4a). These terms include ‘ATP metabolic process’ [114], ‘gluconeogenesis’, ‘mitochondrial calcium ion transport’ [69, 96], ‘glycolytic process’ [131], ‘glycosphingolipid metabolic process’ [19, 58], ‘mitophagy’ [41, 79, 122], ‘positive regulation of mitochondrial fission’ [6, 71], ‘mitochondrial matrix’ [125], ‘mitochondrial membrane’ [25, 125], ‘integral component of mitochondrial inner membrane’ [25], ‘integral component of mitochondrial outer membrane’ [25], ‘protein import into mitochondrial matrix’ [75, 76], ‘negative regulation of amino acid transport’, ‘branched-chain amino acid catabolic process’ [155], ‘peroxisome’ [43, 53], ‘peroxisomal matrix’ [43, 53], all of which point toward altered energy metabolism and mitochondrial dysfunction. Additionally, categories related to cells’ cytoskeletal structure, such as ‘actin filament-based movement’, ‘apical dendrite’ [50], ‘microtubule associated complex’ [50, 147], and ‘anchored component of membrane’ are also implicated in the disease, which point to the structural dysregulations happening in the cells causing axonal transport disturbance [77], impairment of information integration in cells [50], and impairing cell adhesion. A comprehensive list of significant terms can be found in the supplementary material.

**Figure 4.**
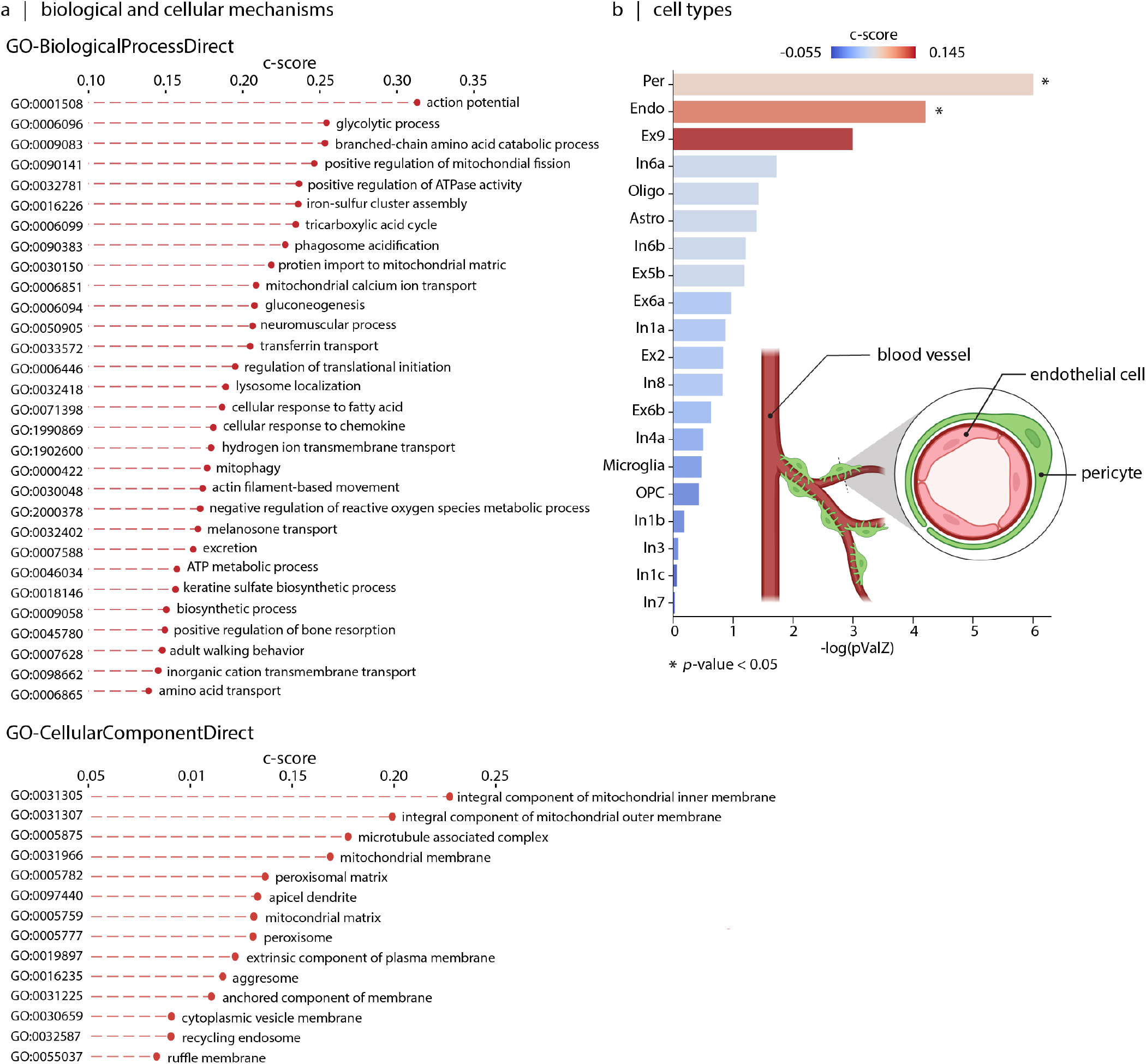
Enrichment analyses of the genes associated with cortical atrophy in ALS. (a) Upper: top 30 biological process terms from the GO Consortium knowledge base associated with gene sets correlated with the ALS epicenter likelihood map [10]. Lower: All cellular components terms from the GO Consortium knowledge base that are significantly enriched in gene sets correlated with the ALS epicenter likelihood map [10]. Terms in both categories are ordered by their category scores (c-scores). (b) Cell type enrichment analysis of ALS epicenter map reveals significant hits for pericytes and endothelial cells [73]. The graphical representation of pericytes and endothelial cells is created with (https://www.biorender.com).

In addition, we estimated cell type enrichment associated with ALS epicenters using genetic markers of different cell types, as determined through single-nucleus droplet-based sequencing and single-cell transposome hypersensitive site sequencing of human brain cells [73] (Fig. 4b). We find significant enrichment for pericytes [31, 110, 150, 152] and endothelial cells [91]. This observation points to possible vascular dysfunction in ALS, potentially explaining the consistent enrichment of metabolic categories. The finding is consistent with previous reports that show a reduction in pericytes in the spinal cord in ALS [46, 65], and interestingly, that these changes occur even prior to the initial clinical manifestations of the disease [128]. Collectively, these findings show that the spatial patterning of atrophy in ALS depends on both network structure as well as local molecular and cellular features that confer greater vulnerability to the disease. In other words, axonal projections are the physical conduit for disease spread, but the trajectory of spreading is guided by local biological features.

### Epicenter location is correlated with clinical presentation and symptom severity

If cortical epicenters reflect the spatial focus of ALS pathology, do they also correlate with the clinical manifestation? To address this question, we analyzed the covariance between individual patient epicenter maps and individual differences across a variety of clinical, cognitive, and demographic variables. We used a multivariate pattern learning algorithm – partial least squares (PLS) – to identify epicenter locations and clinical subtypes that maximally covary with each other [70, 88, 156] (see *Methods*).

The analysis revealed two latent variables that accounted for 27.85% and 13.54% of covariance between epicenter maps and clinical scores (Fig. 5, Fig. S4). Both latent variables were statistically significant using permutation tests (*p* = 9.99×10^−4^, 1.30×10^−2^), but only the first latent variable could be cross-validated (*p* = 0.030). The first latent variable captures a mainly primary motor cortical epicenter pattern. Individuals who display this epicenter pattern tend to have worse motor function, including abnormal index finger and foot tapping scores, daily physical functions (Revised Amyotrophic Lateral Sclerosis Functional Rating Scale; ALSFRS scores), and muscle tone. Interestingly, pathology in these epicenters is uncorrelated with most cognitive scores, except for worse spelling, cube counting, and visuo-spatial scores. In other words, atrophy load in this cortical location is linked with worse motor symptoms and a higher level of disability.

**Figure 5.**
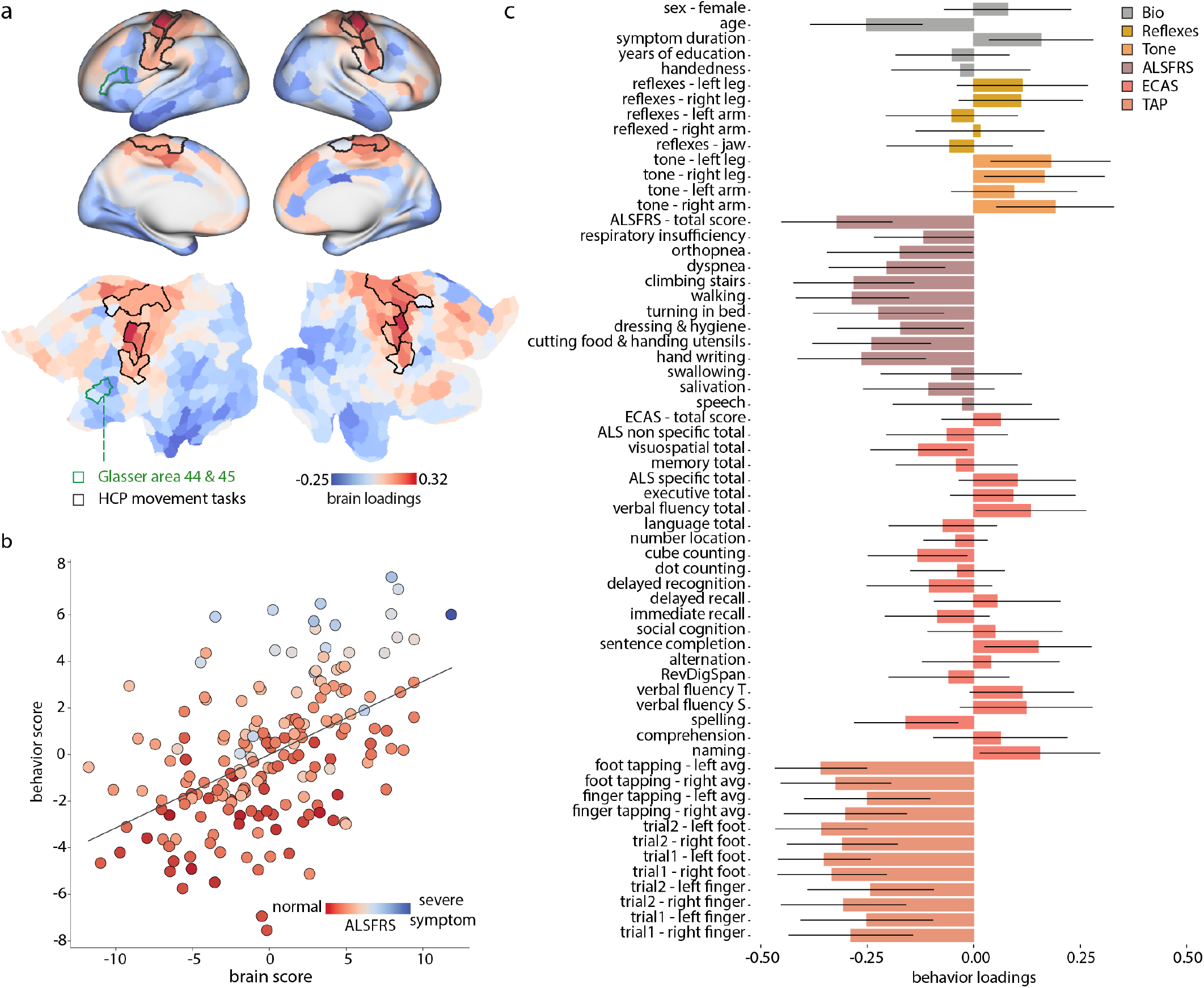
Relating individual epicenter maps with clinical and behavioural measures. (a) The first latent variable from a PLS analysis relating individual epicenter maps and clinical-behavioural measurements. Brain loadings are shown on the fs-LR inflated and flat cortical surfaces. Regions demarcated by the black border are those with the greatest effect sizes (Cohen’s d effect size greater than 1) in the group-average activation map from S1200 Human Connectome Project package for the movement task contrasts [12]. Regions demarcated by green borders showcase areas 44 and 45 from the Glasser parcellation [51]. These regions, specifically in the left hemisphere [105], correspond the Broca’s area [16]. (b) The scatter plot visualizes the individual participants’ brain scores versus behavioral PLS scores (Pearson correlation coefficient, *r* = 0.50; Spearman correlation coefficient, *r* = 0.51); each participant’s score is colored based on the revised-ALSFRS total score. The total ALSFRS score quantifies the degree of functional disability resulting from the disease (greater values correspond to lower disease severity) [27]. (c) The bar plot visualizes the behavioral/clinical measures’ loadings. The contribution (effect size) of individual variables is assessed by bootstrap resampling (1, 000 repetitions).

Despite statistical significance and a large effect size, the second latent variable could not be cross-validated. We therefore relegate the figure to the supplement (Fig. S4) and only describe this latent variable for completeness. The second latent variable captures mainly a dorsolateral prefrontal cortical epicenter pattern. Individuals who display this epicenter pattern tend to have worse ALSFRS speech and worse ALSFRS swallowing scores, as well as respiratory insufficiency (i.e. worse ALSFRS-respiratory insufficiency score). The clinical pattern also captures covariance between atrophy in medial prefrontal cortex and lower scores in multiple cognitive measures. Collectively, these patterns suggest that individual differences in epicenter location are closely linked with clinical presentation. Importantly, the two latent variables are reminiscent of spinal-onset and bulbar-onset ALS clinical subtypes, which we examine in detail next.

### Atrophy epicenters in spinal- and bulbar-onset ALS

So far, we focused the analysis on a common atrophy pattern across the patient sample. However, ALS is heterogeneous, and in clinical practice individuals are often stratified according to the initial body region of onset [106]. Most individuals experience spinal (limb) onset of the disease where the first symptoms appear in the legs and/or hands, while a third of patients report the first symptoms being in bulbar areas [86], citing difficulties in salivation, swallowing, and speaking [59, 72]. The present dataset (CALSNIC) provides stratification for both bulbar-(*N* = 38) and spinal-onset (*N* = 140) ALS subtypes (see *Methods*). There are also rare cases of patients who report respiratory difficulties in the initial stages of the disease [123], but these were not included in the dataset. Neither were patients with mixed spinal- and bulbar-onset, nor those with frontotemporal dementia-onset.

Here we investigate whether cortical network epicenters are related to spinal- and bulbar-onset of the disease. We estimate epicenter probability maps for spinal and bulbar ALS with the two methods presented in the *Epicenters of cortical atrophy* subsection (ranking and the SIR modeling). The two methods yield similar epicenter likelihood maps (Pearson correlation coefficient; *r* = 0.80, *p*_spin_ = 9.99 × 10^−4^) for both disease onset types (Fig. 6a). Importantly, epicenter probability maps in spinal and bulbar ALS are different: in spinal-onset ALS atrophy is mainly confined to primary motor cortex, and paracentral lobule; conversely, in the bulbar-onset ALS atrophy infiltrates areas in lower paracentral gyrus and inferior frontal gyrus. To highlight epicenter differences between these two subtypes, we contrast cortical epicenter maps for individuals with spinal and bulbar ALS using *t*-tests. The obtained *t*-statistic map identifies regions that are more likely to be epicenters in one type compared to the other (Fig.6b). For reference, we use the Human Connectome Project motor task group average effect size maps to delineate and overlay borders of cortical regions associated with movement of the tongue, hands and feet. Consistent with their clinical subtypes, bulbar-onset individuals show cortical disease epicenters in regions linked to tongue movement and in Broca’s area, accounting for increased speech deterioration in the bulbar group compared to the spinal group [80]. Conversely, the spinal-onset individuals predominantly have more likely epicenters in regions associated with movement of feet. Additionally, assessing behavioral measures in CALSNIC dataset show that individuals with the bulbar-onset ALS have severe inability in their speech, salivation, and swallowing, and they have increased jaw and upper limb reflexes compared to the individuals with spinal-onset ALS; meanwhile, those with spinal-onset ALS displayed more significant weakness in writing, walking, and stair climbing abilities. These results show how network epicenters of cortical atrophy align with the clinical manifestation of the disease.

**Figure 6.**
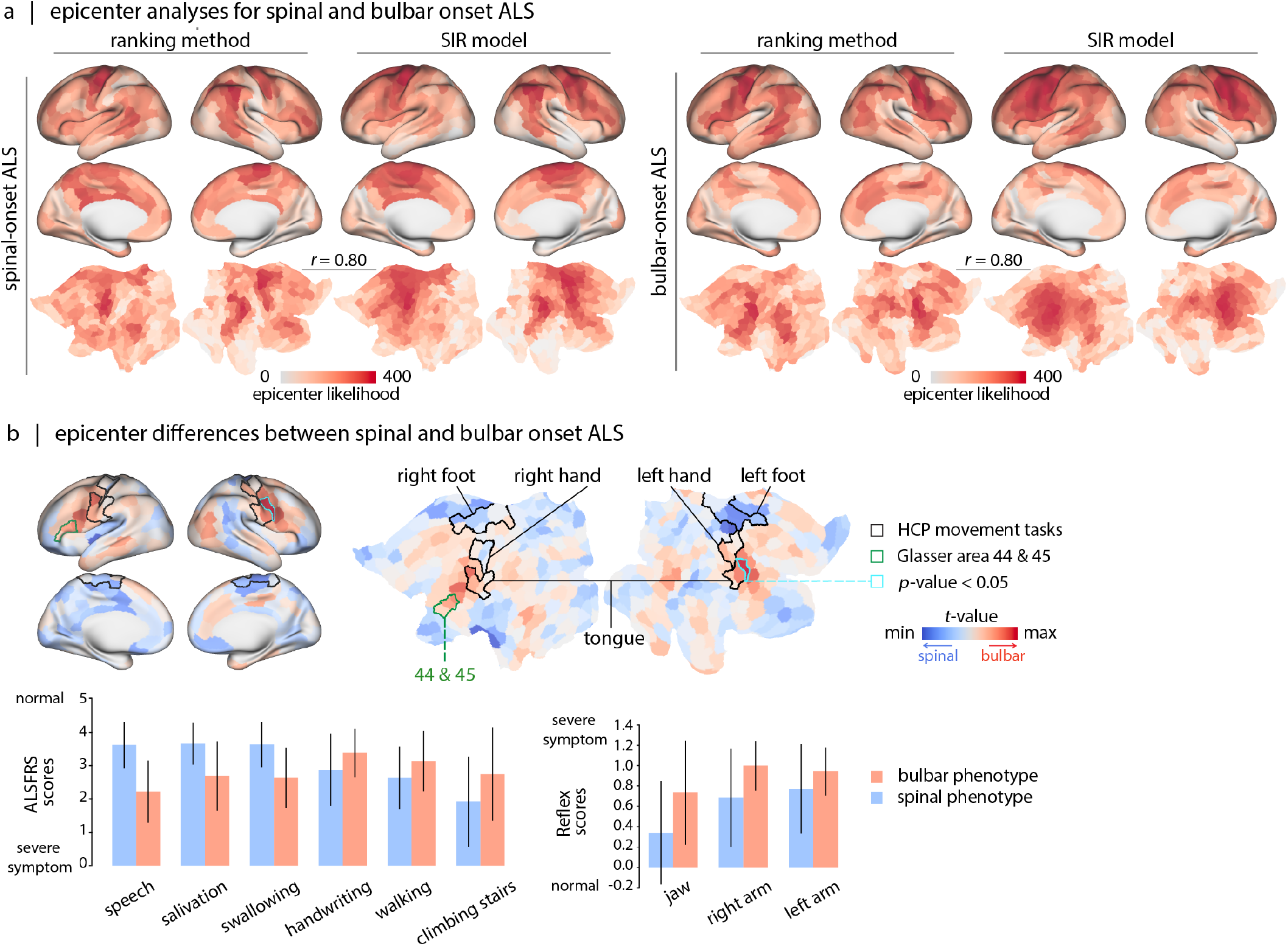
Atrophy epicenters in spinal- and bulbar-onset ALS. (a) Epicenter likelihood maps for spinal-onset ALS and bulbar-onset ALS. Maps are obtained for each subtype using two methodologies: agent-based SIR modeling and a ranking approach. The maps derived by the two methods are correlated with each other, for both subtypes (Pearson correlation coefficient; *r* = 0.80). (b) The brain maps depict the *t*-statistics derived from contrasting the bulbar- versus spinal-onset ALS epicenter maps. Regions demarcated by the black border are those with the greatest effect sizes (Cohen’s d effect size greater than 1) in the group-average activation map from S1200 Human Connectome Project package for the movement task contrasts [12]. Regions demarcated by green borders showcase areas 44 and 45 from the Glasser parcellation [51]. These regions, specifically in the left hemisphere [105], correspond the Broca’s area [16]. Cyan borders highlight two parcels that retain significance following FDR correction. We further contrasted the behavioral/clinical measures for spinal- and bulbar-onset ALS patients. Bar plots show measures that are significantly different between the two subtypes (FDR corrected) and error bars represent the standard deviation. Atrophy maps for each subtype are shown in Fig. S5.

## DISCUSSION

The present report explores how network structure and local biological features jointly shape the spatial patterning of atrophy in ALS. We find that structural connectivity, together with inter-regional similarity of molecular and cellular features, closely aligns with the spatial patterning of atrophy and clinical expression of ALS. We identify consistent and prominent disease epicenters in motor cortex. Epicenter probability maps show transcriptomic enrichment for biological pathways involved in mitochondrial function as well as support cells, such as endothelial cells and pericytes. Finally, individual differences in epicenter location correspond to individual differences in clinical and cognitive symptoms and differentiate patient subtypes.

We consistently find that atrophy in a brain region is correlated with the atrophy of anatomically connected regions, suggesting that atrophy spreads via white matter projections. This is consistent with the notion that ALS pathology is related to misfolding and network spreading of TAR-DNA binding protein (TDP)-43 [26, 115, 148]. According to this account, pathological changes in conformation (misfolding) of endogenous TDP-43 induce aggregation and further misfolding. Trans-synaptic spread of pathogenic misfolded TDP-43 results in patterns of cell death and atrophy that ultimately resemble brain network architecture. Our results support this account in two ways. First, the corticospinal tract exhibits the most pronounced atrophy (consistent with findings in [36, 40, 42, 118]), as does primary motor cortex, suggesting that cortical regions with strong physical connections to the corticospinal tract experience the greatest pathology. Second, a region’s atrophy and the atrophy of its neighbours are correlated, suggesting that network-based exposure to the pathology confers increased risk of pathology. This correlation is significantly greater than in rewired null models that preserve degree sequences and edge lengths, suggesting that the effect is not trivially due to spatial proximity or the total number of connections in a given region, but rather due to the arrangement of connections and the overall topology of the network. Altogether, these results emphasize the role of structural network architecture as the mediator of the pathological protein spread in ALS [89, 98, 112, 142].

In addition to structural connectivity, we find that the local biological features of brain regions contribute to disease spread. Numerous theories posit that the propagation of ALS may also be due to genetic mutations [28, 34, 67, 68] and atypical metabolic function [38, 137]. Consistent with this work, we find that other types of inter-regional similarity, such as gene expression, neurotransmitter receptor, laminar, metabolic, and hemodynamic similarity also contribute to the patterning of atrophy. Augmenting white matter connections with connection weights from these similarity matrices, we observe increased correlations between regional atrophy and atrophy of connected regions. In other words, pathology is mediated by structural connectivity, but spreading is more likely to take place between neuronal populations that share similar molecular and cellular characteristics.

Given that disease spread appears to follow network structure, we next sought to identify the most likely cortical epicenters. We assessed the likelihood of various brain regions acting as cortical disease epicenters using two methods: (1) data-driven ranking, and (2) agent-based SIR modeling. Both methods identified regions located in the primary motor area to be the likeliest epicenters, as well as regions in frontal and temporal cortex, such as the temporoparietal junction. These results are in line with histopathological findings, showing TDP-43 pathology in motor cortex (Brodmann areas 4 and 6) in the initial stages of the disease [20]. The involvement of frontal and temporal areas in later stages of the disease is also well-documented [20, 36, 143].

What biological features predispose regions to act as disease epicenters? The epicenter probability map is correlated with the spatial expression of genes involved in metabolic pathways, pointing toward mitochondrial functions such as ATP production, as well as the structural organization of mitochondrial membrane and matrix, and disruption of mitochondrial calcium ion transport. This finding is in line with the idea of mitochondrial stress in ALS. Namely, dysregulation of mitochondrial calcium transport, potentially driven by increased excitatory cytosolic calcium uptake or changes in the function of mitochondrial calcium transporters, results in mitochondrial injury and triggers mitophagy. In addition, mitochondrial dysfunction is thought to cause further structural abnormalities in multiple cell types; for example, deletion of mitochondrial calcium uniporter changes dendritic spine morphometry [100].

In addition, the epicenter likelihood map overlaps with transcriptomic signatures of pericytes and endothelial cells. Endothelial cells are single-layer cells lining the blood vessels [64]. Pericytes surround the endothelial cells by extending their long cytoplasmic processes on the surface of endothelial tubes. The interaction between endothelial cells and pericytes is needed in formation and maintenance of blood-brain barrier (BBB), and blood-spinal cord barrier (BSCB); which contribute to maintaining the controlled chemical composition of the neuronal milieu [159, 160]. Multiple studies point to a breakdown in both BBB and BSCB in ALS [32, 151]. In mouse models of ALS, damage to BSCB, BBB, and endothelial cells are observed early in the disease [45, 48], and often precede the onset of motor symptoms [158]. Similar vascular deficits have also been reported in human patients [92]. Indeed, cerebrospinal fluid abnormalities in ALS are thought to at least partly originate from the increase in BBB permeability [8, 74]. Postmortem histology confirms the loss of integrity of BBB and SCSB due to endothelial cell damage and pericyte degeneration [46, 150]. Our findings corroborate that the scope of inquiry for studying ALS should be broadened beyond motor neurons, and that support cells – such as vascular cells – should also be considered as therapeutic targets. Interestingly, recent findings suggest that treatment of ALS mice models with unmodified human bone marrow CD34+ (hBM34+) cells accelerates BSCB repair, leading to differentiation into endothelial cells, reduced astrogliosis and microgliosis, and improved perivascular integration, ultimately promoting survival of motor neurons [47]; likewise, treatment with pericytes improves overall survival in SOD1 mutant ALS mice [33].

Finally, the locations of individual network epicenters map onto individual differences in clinical manifestation. Overall, individuals with greater epicenter probability in motor cortex had worse motor symptoms and signs, including poorer finger and foot tapping, increased muscle tone and reflexes, and lesser capability to perform daily functions involving the limbs. The multivariate clinical subtype also included lower scores in naming, sentence completion, and verbal fluency tasks, presumably corresponding to greater epicenter probability in left inferior frontal cortex (Broca’s area; Fig. 5). Interestingly, disease epicenters in patients with bulbar- and spinal-onset of the disease evolve in different trajectories. Specifically, patients with bulbar-onset ALS displayed cortical epicenters located in areas that overlapped with parts of somatomotor area involved in tongue movement. In contrast, the likeliest epicenters for the spinal-onset patients were near areas involved in limb movements. Collectively, these results demonstrate that studying ALS pathology from a network perspective can help to trace the spatial origin of the disease, to identify the molecular and cellular contributions to pathology, and, ultimately, to map individual differences in brain-behaviour relationships and clinical subtypes.

The present results should be considered with respect to several methodological limitations. First, we estimated atrophy using *in vivo* MRI deformation-based morphometry, a technique that is well-validated but that does not directly measure pathology (e.g. neuronal loss, TDP-43 deposition). Despite this limitation, the results recapitulate numerous histopathological hallmarks of the disease. Second, all network spreading effects were estimated using an independent high-resolution diffusion MRI dataset, rather than individual patient connectomes. This methodological decision was made to compensate for potential inaccuracies in individual-level diffusion tractometry, and highlights the need for more multimodal imaging in patient samples. Likewise, inter-regional transcriptomic similarity, receptor similarity, laminar similarity, metabolic similarity and hemodynamic similarity matrices are estimated from non-ALS participants. Third, all results are based on a single dataset. Although we used multiple methods whenever possible (e.g. when identifying epicenters), as well as cross-validation (e.g. for the PLS model), ideally the results should be independently confirmed using a different ALS dataset.

In conclusion, we reveal that network structure and local biological features leave an indelible mark on the course and expression of neurodegeneration in ALS. These two factors are closely related and should be studied simultaneously, rather than in isolation. Conceptualizing neurodegeneration as a multiscale spreading process may help to identify molecular, cellular and regional targets for therapies that slow or divert the course of pathology.

## METHODS

All codes used to perform the analyses are available at https://github.com/netneurolab/Farahani_ALS.

### CALSNIC dataset

Data were retrieved from the Canadian ALS Neuroimaging Consortium (CALSNIC) dataset (http://calsnic.org). The dataset comprises data from individuals diagnosed with possible, probable, or definite ALS, according to the revised El Escorial Criteria [22], alongside data from age-matched healthy controls. All participants underwent magnetic resonance imaging (MRI), yielding 1 mm isotropic T1-weighted (T1w) data for each individual. We use data acquired from eight different imaging sites, including: University of Calgary (CAL), University of Alberta (EDM), McGill University (MON), University of Toronto (TOR), University of British Columbia (VAN), University of Miami (MIA), Université Laval (QUE), and University of Utah (UTA). We exclude data from participants if they had other CNS abnormalities or reported psychiatric illness. The CALSNIC dataset provides longitudinal neuroimaging data with approximately four-month intervals between sessions for some but not all participants. We only keep individuals with a scan acquired at baseline for further analysis, resulting in 192 individuals with ALS (70 female; mean age: 59.62 ± 10.20) and 175 healthy control participants (96 female; mean age: 55.41 ± 10.00). Note that data used in this study comes from two phases of the CALSNIC dataset launched to date, which have slight changes in the structural MR imaging paradigm. Consequently, when building the ordinary least square model to estimate disease-related atrophy, if data from the same imaging site is acquired during different project phases, we treat the data from each phase as a separate entity. This assumption leads to considering twelve imaging sites in total when building the model (CAL: phase 1, 2; EDM: phase 1, 2; MON: phase 1, 2; TOR: phase 1, 2; VAN: phase 1; MIA: phase 2; QUE: phase 2; UTA: phase 2). The detailed data description is provided in the original publication [66].

### Behavioral/clinical measures

The CALSNIC dataset, in addition to the magnetic resonance imaging data, provides behavioral/clinical assessments for individuals. These encompass ALS-related motor and cognitive evaluations, including Revised Amyotrophic Lateral Sclerosis Functional Rating Scale (ALS-FRS), Edinburgh Cognitive and Behavioural ALS Screen (ECAS), finger/foot tapping test, and evaluations for abnormal muscle tone and reflexes. Data on individuals’ sex, age, symptom duration, years of education, and handedness is also provided. Below is a brief overview of each measure.

ALSFRS is a rating measure that quantifies the level of disability in ALS patients. Higher values of this scale (with a maximum of 48), indicate lower disease severity. Specifically, ALSFRS assesses the participants’ functional capability in speech, salivation, swallowing, hand-writing, gastrostomy, cutting food and handling utensils, dressing and hygiene, turning in bed and adjusting bed clothes, walking, climbing stairs, dyspnea, orthopnea, and respiratory insufficiency [27]. ECAS is a cognitive screening assessment measuring executive function, letter and semantic fluency, attention, memory, language, and visuospatial function of individuals [2]. A higher total score for ECAS (up to 136) signifies better cognitive performance. The tapping score, which is a clinical motor symptom severity indicator, quantifies the number of taps a participant can perform in 10 seconds by their fingers or feet. A higher count in this test reflects better motor function. In CALSNIC dataset, the tapping scores for both index-finger, and foot are measured two times per participant. Muscle tone and reflex measures are also included, with lower scores signifying normal tone and reflex conditions.

In this study, we use behavioral/clinical measures to relate the cortical epicenter locations to the behavioral manifestations of individuals with ALS. Of the 192 patients included in the study, 8 individuals have incorrect ECAS administration; these participants are therefore excluded from analyses relating brain and behavior data.

### Deformation based morphometry

T1w data are preprocessed and Deformation-Based Morphometry (DBM) maps are derived per participant using the Montreal Neurological Institute Medical Imaging NetCD (MNI-MINC) tools, publicly available at https://github.com/BIC-MNI/minc-tools. The pre-processing steps include image denoising [35], intensity inhomogeneity correction [124], and image intensity normalization into range (0–100) using histogram matching; next, each T1w image is first linearly and then nonlinearly registered into the MNI152-NonLinear2009cSym standard brain. DBM maps are derived by estimating the local deformation needed in each voxel in an individual’s T1w image to nonlinearly match it to the standard template. The required deformation is estimated by the Jacobian determinant of the inverse nonlinear deformation field and can be used as an indirect estimate of brain atrophy [29, 49]. DBM values lower than 1 indicate that the corresponding region is smaller in the participant than in the template (atrophy in the participant compared to the template). Conversely, DBM values greater than 1 indicate that the corresponding region in the participant space is larger than the same region in the template space (expansion in the participant compared to the template). Collectively, the DBM maps encode the morphological differences between the T1w data of a given participant and the standard brain defined by the MNI template.

### Atrophy maps

The Jacobian determinant from the DBM analysis serves as a dependent variable, influenced not only by the diagnosis but also by factors such as age, sex, and imaging site. To isolate factors unrelated to diagnosis from the DBM maps of ALS patients and to obtain a more accurate measure of disease-specific atrophy, we compute *w*-score maps per individual [61]. The *w*-score value at each voxel quantifies the normalized deviation of the observed DBM value from its expected DBM value, adjusted for age, sex, and imaging site. The expected value for DBM at each voxel is estimated using an ordinary least squares model constructed based on control participants’ data. Greater absolute disparity between the observed and expected value of DBM indicates more severe atrophy or expansion in a given voxel. The formula to calculate the *w*-score is provided in the following:

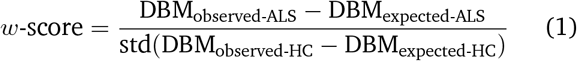

where *std* stands for standard deviation. A negative *w*-score in this context signifies greater atrophy than the mean value expected for a healthy participant. After calculating the *w*-score maps of all ALS participants under study, *w*-score maps are averaged across ALS patients to create a single collective atrophy map. To simplify interpretation, the *w*-score maps are multiplied by −1 so that the larger numerical values correspond to more atrophy.

Throughout the manuscript, the *w*-score maps (post-inversion) will be referred to as the “atrophy map”. The group-average ALS atrophy map is visualized in Fig. 1a.

### Anatomical parcellation

ALS atrophy maps are parcellated using two different atlases. For cortical regions, we apply the Schaefer-400 parcellation [111], and to examine the sub-cortex, specifically the white matter tracts, we use the Johns Hopkins University (JHU) white matter tractography atlas (thresholded at 25% probability) [60, 146].

### Network reconstruction

Brain networks (structural connectivity and inter-regional similarity networks) are retrieved from netneurolab [13, 56]. Below we briefly describe how each network is reconstructed.

#### 1 Structural connectome

Structural connectivity is a matrix that provides information regarding the white matter connections across pairs of brain regions. In this study, the dataset used for building the connectome comes from 327 unrelated healthy participants (145 males; age: 22–35) in the Human Connectome Project (HCP-S900 release), scanned using a 3T Connectome Skyra scanner. Diffusion MRI data is preprocessed using the HCP minimal preprocessing pipeline. For detailed acquisition information and preprocessing steps refer to these references [52, 136]. Structural connectome is reconstructed from the diffusion MRI using the MRtrix3 package (https://www.mrtrix.org) [133]. Grey matter is parcellated based on the Schaefer-400 atlas, and fiber orientation distributions are generated using a multi-shell multi-tissue constrained spherical deconvolution algorithm [63, 111]. Initially, a tractogram is generated with 40 million streamlines, with a maximum tract length of 250 and a fractional anisotropy cut-off of 0.06. Spherical-deconvolution informed filtering of tractograms (SIFT2) is used to reconstruct whole brain streamlines weighted by cross-section multipliers [126]. For further insights into individual network reconstructions, please consult the reference provided [103]. A group consensus structural network is then created such that the mean density and edge length distribution observed across individual participants is preserved [15]. The weights of the edges in the consensus networks represent the log-transform of the number of streamlines in the parcels, averaged across participants for whom these are present [13].

#### 2 Hemodynamic similarity

Hemodynamic similarity, often referred to as functional connectivity, summarizes the similarity across brain regions in terms of the synchronization and similarity of their co-fluctuation in the BOLD signal. The data incorporated to build the connectome comes from 326 unrelated healthy participants (145 males; age: 22–35) in the Human Connectome Project (HCP-S900 release), scanned using a 3T Connectome Skyra scanner [136]. In this dataset, each participant has undergone four 15-minute resting-state functional MRI scans, each with a TR of 720 ms. Data is preprocessed using the HCP minimal preprocessing pipeline. For detailed preprocessing steps, refer to the cited reference [52]. The voxel-wise functional MRI data is parcellated using the Schaefer-400 atlas [111]. The parcellated time-series are then used to construct functional connectivity matrices, computed as Pearson correlation coefficient between pairs of regional time-series for each of the four scans per participant. To obtain a group-level functional connectivity matrix, mean functional connectivity across all participants and scans is computed. This matrix is normalized using Fisher’s r-to-z transformation [56].

#### 3 Metabolic similarity

Metabolic similarity estimates the similarity between brain regions in terms of glucose metabolism or, in other words, in terms of energy consumption. This network is reconstructed using positron emission tomography (PET) images of the [F18]-fluordoxyglucose tracer. The dataset includes 26 healthy participants (77% female; age: 18–23) who participated in a 95-minute simultaneous MR-PET scan acquired using a 3T molecular MR scanner [62]. PET images are preprocessed according to [144]. Each volume of the PET time-series is registered to the MNI152 template space and is parcellated according to the Schaefer-400 atlas [111]. Parcellated time-series from pairs of brain regions are then correlated (Pearson’s correlation coefficient) to construct a metabolic connectivity matrix for each participant. Subsequently, by averaging connectivity matrices across all participants, a group-average metabolic connectome is obtained. This matrix is normalized using Fisher’s r-to-z transformation [56].

#### 4 Gene expression similarity

Correlated gene expression quantifies the transcriptomic similarity between pairs of brain regions. The underlying data to construct this connectome comes from the bulk tissue microarray expression data collected from six post-mortem brains (1 female; age: 24–57, mean age: 42.50±13.38).

This data is provided by the Allen Human Brain Atlas (https://human.brain-map.org) [57] and is processed using the abagen toolbox, publicly available at https://github.com/rmarkello/abagen [82], yielding a map for each gene in the parcellated MNI template (Schaefer-400 [111]). Genes with high differential stability across donors (threshold of 0.1) are considered for the analysis, resulting in 8, 687 stable genes. A region×region correlated gene expression matrix is constructed by correlating normalized gene expression profiles between pairs of brain regions (Pearson correlation coefficient). This matrix is then normalized using Fisher’s r-to-z transformation for subsequent analysis [56].

#### 5. Receptor similarity

Receptor similarity measures how correlated the receptor density profiles are between brain regions. To construct this network, PET tracer images for 18 neurotransmitter receptors and transporters are used [54, 83]. These receptors/transporters cover nine neurotransmitter systems, including dopamine (D1, D2, DAT), norepinephrine (NET), serotonin (5-HT1A, 5-HT1B, 5-HT2, 5-HT4, 5-HT6, 5-HTT), acetylcholine (*α*4*β*2, M1, VAChT), glutamate (mGluR5), GABA (GABAA), histamine (H3), cannabinoid (CB1), and mu-opioid (MOR). Each of these PET tracer images is parcellated based on the Schaefer-400 atlas [111] and normalized using z-scores. A region×region receptor similarity matrix is constructed by correlating receptor profiles across all pairs of brain regions (Pearson correlation coefficient). This matrix is then normalized using Fisher’s r-to-z transform [56].

#### 6. Laminar similarity

Laminar similarity, estimated from histological data, assesses the similarity in cellular distributions across cortical layers within pairs of brain regions [56, 102]. Data were recovered from the high-resolution (20 μm) BigBrain atlas, a postmortem Merker-stained histological atlas of a 65-year-old male [7]. Staining intensity profiles are sampled across 50 equi-volumetric surfaces within the cortical grey matter, enabling the assessment of neuronal density and soma size variations across cortical layers. These intensity profiles also help delineate boundaries among cortical layers, such as supragranular (layers I–III), granular (layer IV), and infragranular (layers V–VI). The BigBrainWarp toolbox [101] was used to transform the data to the surface-based fs-LR template, which is then parcellated based on the Schaefer-400 atlas [111]. A laminar similarity matrix is estimated by computing the partial correlation between regional intensity profiles. The matrix is then normalized using Fisher’s r-to-z transform [56].

### Disease exposure

In this section, our objective is to assess how a specific biological brain connectome affects the spread of pathology across the cortex in ALS. We initially apply a threshold to the connectome under study (e.g. metabolic similarity) to retain only positive values; the thresholded connectome is considered as a “network”, whose “nodes” correspond to the Schaefer-400 parcels [111] and the “edges” are defined based on the values in the off-diagonal elements of the connectivity (similarity) matrix. We assume that these edges provide potential pathways for pathology propagation. At each node, we define the global disease “exposure” as a measure quantifying the extent of atrophy its neighbouring nodes are experiencing, weighted according to the strength of the edges that connect the node with its neighbours. Here, “neighbours” are defined as nodes directly connected to the node in focus.The following is a mathematical formulation of disease exposure at a given node, denoted as node *i*:

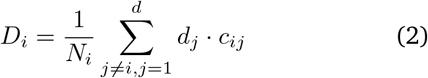

Here *D*_*i*_ represents the disease exposure at node *i*; which is calculated as the weighted average of atrophy across the neighbours of node *i*. The atrophy values of neighbours are denoted by *d*_*j*_, where *j* is the neighbour’s identifier. Each neighbour atrophy value, *d*_*j*_, is multiplied by the edge strength between region *i* and node *j*, represented by *c*_*ij*_. *N*_*i*_ in the formula represents the number of connections made by node *i* within the network. To assess whether connectome architecture is related to the spatial patterning of atrophy, we correlate the degree of atrophy in individual nodes and their respective disease exposures. This analysis is repeated while considering various brain connectomes (including structural connectivity, gene-expression similarity, receptor similarity, laminar similarity, metabolic similarity, and hemodynamic similarity).

### Epicenter mapping: data-driven method

With an atrophy map and an underlying connectome, it is possible to identify putative disease epicenters using either epidemiological data-driven methods, or computational modeling [119, 120]. The data-driven approach is based on the notion that one of the main factors of disease spread is the existence of structural connectivity across brain regions (nodes). Here an epicenter is defined as a node that is both highly atrophied, and physically connected to nodes that are also highly atrophied (Fig. 2b). Epicenters are identified using two separate ranking procedures. In the first ranking procedure, nodes are ranked in ascending order according to their mean atrophy. The second ranking procedure is performed on the array containing the weighted neighbour atrophy values (nodes’ exposure values). The disease exposure value at each node is calculated as previously described in *Disease Exposure*. This new set of values per node is also ranked in ascending order, quantifying the involvement of node’s neighbours in pathology. Finally, we calculate the average ranking of a node in the two lists, reflecting its likelihood as a disease epicenter[23, 55, 99, 116, 119, 120].

### Epicenter mapping: agent-based spreading model

Disease epicenters can also be identified using a modeling approach known as the agent-based Susceptible-Infected-Removed (SIR) model [157]. In brief, this model simulates the brain spread of pathology considering the structural connectome as a network through which misfolded proteins (agents) can propagate [21]. In the case of ALS, these agents may represent TDP-43 [26, 115, 148].

We apply this methodology to find epicenters (model parameters outlined in Table S1). The epicenters are identified as the nodes that—if chosen as the disease’s initial point of pathology (seed region)—will lead to the highest Pearson correlation coefficient value between the simulated and observed cortical atrophy pattern during the spread of the agent. Here, the disease spread simulation is performed over a total of 10, 000 time steps. Nodes are then ranked based on the maximum correlation value between the observed patterns of cortical atrophy and the simulated atrophy patterns when each node is used as the initial seed.

### Gene category enrichment analysis

Biological pathways that are correlated with the ALS epicenter likelihood map were identified using a gene category enrichment analysis (GCEA). Cortical maps for biological pathways were defined according to the gene expression data coming from the Allen Human Brain Atlas [57]. The data was preprocessed and mapped to parcellated brain regions using the abagen toolbox, publicly available at https://github.com/rmarkello/abagen [82]. To perform the enrichment analysis, we use the ABAnnotate Matlab-based toolbox, publicly available at https://github.com/LeonDLotter/ABAnnotate [78]. The package is adapted from the toolbox developed by Fulcher and colleagues https://github.com/benfulcher/ GeneCategoryEnrichmentAnalysis [44]. The GCEA procedure assesses whether genes in a particular category are more correlated with a given brain phenotype than a random phenotype with comparable spatial autocorrelation (ensemble-based null model) [44].

To address spatial auto-correlation effects, 30, 000 spatially auto-correlated null maps are generated from the epicenter likelihood map using the neuromaps toolbox (method = “vasa” [83, 139]) and are inputted to ABAnnotate package for the testing procedure. After matching category and Allen Human Brain genes based on gene symbols, and removing the genes with differential stability lower than 0.1, the Pearson correlations between the epicenter map, the null maps, and all gene expression maps are calculated. For each null map and each category, null category scores are obtained as the mean z-transformed correlation coefficients. Positive-sided *p*-values, indicative of the relationship between the epicenter map and each category, are determined by comparing the actual category scores to the null distribution, with subsequent False Discovery Rate (FDR) correction applied. For gene-category annotations, we use the GO biological and cellular processes [10] as well as the cell-type categories introduced by [73].

### Null models

Reported associations among brain maps and/or networks are assessed with respect to three null models [138]. Here we briefly outline the logic and implementation of each. First, to assess the effect of spatial autocorrelation on spatial associations between brain maps, we use the so-called spatial auto-correlation preserving permutation tests, commonly referred to as “spin tests” [5]. Briefly brain phenotypes (e.g. cortical atrophy maps) are projected to spherical projection of the fsaverage surface. This involves selecting the coordinates of the vertex closest to the center of mass for each parcel. These parcel coordinates are then randomly rotated, and original parcels are reassigned to the value of the closest rotated parcel (*n* repetitions). For parcels where the medial wall is the closest, we assign the value of the next closest parcel instead. Following these steps, we obtain a series of randomized brain maps that have the same values and spatial autocorrelation as the original map but where the relationship between values and their spatial location has been permuted. These maps are then used to generate null distributions of desired statistics, such as null node-neighbour correlation values [141].

When evaluating the role of the structural connectome in disease spread, we use two additional network randomization methods [138]. One approach constructs degree-preserving randomized networks [85, 138]. In this case, the atrophy map is unchanged, but the structural connectome itself is randomized 1, 000 times. In each randomization realization, each edge is rewired 10 times to generate a randomized network with the same size, density and degree sequence as the actual network [85]. Using these null networks, the Pearson correlation coefficient between node and its neighbours’ is recalculated, and a two-sided *p*-value is estimated in a non-parametric manner.

Finally, the most conservative approach to test the role of structural connectome in disease spread involves using degree- and edge length-preserving randomized networks. In this approach, the atrophy map is kept unchanged, but the structural connectome is randomized 1, 000 times to create randomized networks with preserved size, density, degree sequence and edge length (sometimes referred to as “cost”) [14, 138]. To achieve this, edges within the structural connectome are categorized based on Euclidean distance into 10 bins. Within each bin, pairs of edges are selected randomly and swapped, with the total number of swaps equaling the number of regions in the network multiplied by 20. This process is repeated 1, 000 times, yielding 1, 000 randomized structural networks. These randomized networks are then used to construct a null distribution for the node-neighbour Pearson correlation coefficient, and a two-sided *p*-value.

### Partial least squares

The goal of partial least squares (PLS) analysis is to relate two data matrices to each other [87, 88]. In the present case, the two matrices represent epicenter likelihood maps (participants×regions) and clinical-behavioural data (participants×measures). The analysis is initialized by computing the covariance between brain (**X**) and behaviour features (**Y**). The resulting covariance matrix is subjected to singular value decomposition:

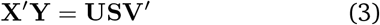

where **U** and **V** are orthonormal matrices of left and right singular vectors and **S** is a diagonal matrix of singular values. Each combination of a left singular vector, a right singular vector, and a singular value constitutes a latent variable. The elements of each singular vector weight the contribution of individual features to the overall multivariate pattern. In the present analysis, these weights correspond to a spatial pattern of cortical epicenters and clinical phenotypes that optimally covary with each other. To estimate the extent to which individual patients express these atrophy or behavioural patterns, patient-specific brain and behavioural scores are calculated. Scores are computed by projecting the original data onto the respective singular vector weights, such that each individual is assigned a brain and a behavioral score, indicating the degree to which a patient expresses each atrophy pattern and behavioural phenotype [88].

The proportion of covariance accounted for by each latent variable is a measure of effect size and is quantified as the ratio of the squared singular value to the sum of all squared singular values. The statistical significance of each latent variable is estimated by permutation testing.

This involves randomly permuting the order of observations (i.e. rows) of data matrix **X** for a total of 1, 000 repetitions, followed by constructing a set of “null” brain-behavior correlation matrices for the permuted brain and unchanged clinical data matrices. These “null” correlation matrices are then subjected to SVD, to generate a distribution of singular values under the null hypothesis that there is no association between brain epicenters and behavioral measures. A non-parametric *p*-value can be estimated for a given latent variable as the probability that a permuted singular value exceeds the original, non-permuted singular value.

The contribution of individual features to the model is estimated using bootstrap resampling. Participants (rows of data matrices **X** and **Y** are randomly sampled with replacement (1, 000 repetitions), resulting in resampled correlation matrices that are then subjected to SVD. This bootstrapping procedure generates a sampling distribution for each singular vector weight. A bootstrap ratio for each behavioral measure is then computed, defined as the ratio of its singular vector weight and its bootstrap-estimated standard error. High bootstrap ratios are indicative of features that make a large contribution to the latent variable and are consistent across samplings.

Finally, we use cross-validation to evaluate the out-of-sample correlation between cortical epicenter patterns and behavioral features. We use 100 random divisions of the dataset, allocating half of the data for training and the other half for testing. In each repetition, we apply PLS to the training data and estimate singular vector weights. Subsequently, each realization of the test data is projected onto the derived weights derived from the training set. We then estimate patient-specific scores and their correlation in the test sample. The procedure is repeated 100 times to establish a distribution of out-of-sample correlation values. To assess the statistical significance of these out-of-sample correlation values, we conduct permutation tests (100 repetitions). During each permutation, we shuffle the epicenter matrix rows and repeat the analysis to create a null distribution of correlation coefficients between epicenter and clinical scores in the test sample. This null distribution is then used to estimate a non-parametric *p*-value.

### Somatotopic map from the HCP movement task

To localize the cortical activation boundaries associated with different body part movements, we utilize the Human Connectome Project’s group-average activation maps from the HCP-S1200 release. The group-average activation maps include the average strength of functional activation across 997 healthy young adults (532 female; age: 22–35) who completed 3T task fMRI runs (for detailed information on fMRI acquisition parameters and the preprocessing steps, refer to [12, 52, 135]). Here, we incorporate a set of group-average motor task activation maps, which reveal the somatotopic organization of the sensorimotor cortex. These motor tasks are originally developed by Buckner et al. [24] and Yeo et al. [154] and involve participants responding to visual prompts to execute specific movements. These movements include finger tapping (left and right), toe squeezing (left and right), and tongue moving. Each movement type is performed in a 12-second block, encompassing 10 movements, and is preceded by a 3 second cue. There are two runs per participant in total, each containing 13 blocks: two for tongue movements, four for hand movements (split evenly between right and left), four for foot movements (also evenly split), and three 15-second fixation blocks. We parcellate the Cohen’s d group-average maps according to the Schaefer-400 atlas [111]. The parcellated map is then thresholded to highlight regions with significant effect sizes (Cohen’s d greater than 1), enabling us to associate the cortical parcels with specific motor functions.

### Contrasting spinal- and bulbar-onset ALS

To compare cortical epicenter likelihood maps between spinal- and bulbar-onset ALS subtypes, we conduct a *t*-test for each cortical brain parcel. This analysis allows us to identify cortical areas that differ across the two disease subtypes in terms of their capacity to spread the disease. Here, individuals are assigned to the spinal subtype (*N* = 140) when the onset region is specified as either “upper motor neurons”, “lower motor neurons”, or a combination of both. Conversely, individuals reporting an onset region of “bulbar”, “bulbar-speech”, or “bulbar-speech and bulbar-swallowing” are categorized under the bulbar ALS subtype (*N* = 38). We use individual epicenter likelihood maps as inputs for the *t*-test models, resulting in a map of *t*-statistics that illustrates the cortical epicenter differences between ALS subtypes. To ensure the reliability of the differences in epicenter locations between the two groups, we apply FDR correction to the *p*-values from the *t*-tests (*n* = 400). This correction signifies two cortical parcels in the right motor area, involved in the tongue movement, as the regions that are statistically different across the subtypes in terms of their epicenter likelihood.

We also examine the behavioral/clinical differences across the disease subtypes. We compare groups (bulbar- and spinal-onset ALS) using *t*-tests for each metric, followed by adjustment for multiple comparisons using FDR. 64 different metrics are included for assessment in total, after removing the metrics which are not reported for more than 20 participants. If missing data is available, it is imputed with the median of the measure. We find significant differences in metrics including “ALSFRS-Speech” (FDR corrected, *p* = 4.21 × 10^−9^); “ALSFRS-Salivation” (FDR corrected, *p* = 4.91 × 10^−5^); “ALSFRS-Swallowing” (FDR corrected, *p* = 4.28 × 10^−6^); “ALSFRS-Handwriting” (FDR corrected, *p* = 0.012); “ALSFRS-Walking” (FDR corrected, *p* = 0.028); “ALSFRS-Climbingstairs” (FDR corrected, *p* = 0.020); “Reflexes-Jaw” (FDR corrected, *p* = 2.46 × 10^−3^); “Reflexes-RightArm” (FDR corrected, *p* = 1.09 × 10^−5^); “Reflexes-LeftArm” (FDR corrected, *p* = 0.02); and “Symptom-Duration” (FDR corrected, *p* = 0.023) (Fig. 6b).

## Acknowledgments

We thank Filip Milisav, Andrea Luppi, Zhen-Qi Liu, Eric G. Ceballos, Moohebat Pourmajidian and Alicja Monaghan for their comments and suggestions on the manuscript. BM acknowledges support from the Natural Sciences and Engineering Research Council of Canada (NSERC), Canadian Institutes of Health Research (CIHR), Brain Canada Foundation Future Leaders Fund, the Canada Research Chairs Program, the Michael J. Fox Foundation, and the Healthy Brains for Healthy Lives initiative. The Canadian ALS Neuroimaging Consortium (CALSNIC) is supported by grants from CIHR, ALS Society of Canada, Brain Canada Foundation, and the Shelly Mrkonjic Research Fund.

---

**Figure S1.**
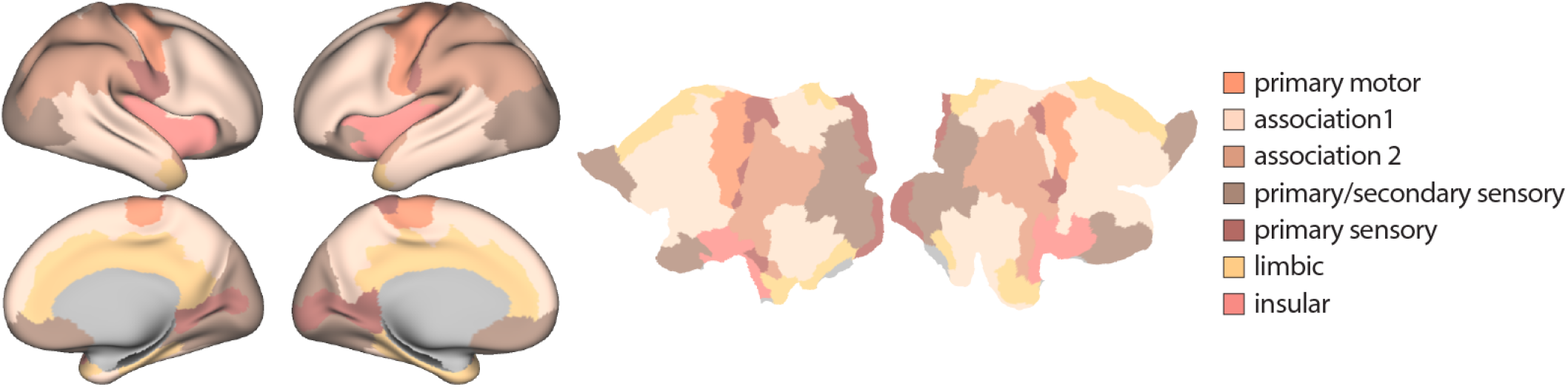
Von Economo’s cytoarchitectonic classes. Von Economo cytoarchitectonic parcellation [39, 113, 145] is shown on the fs-LR inflated (left) and flat (right) cortical surfaces.

**Figure S2.**
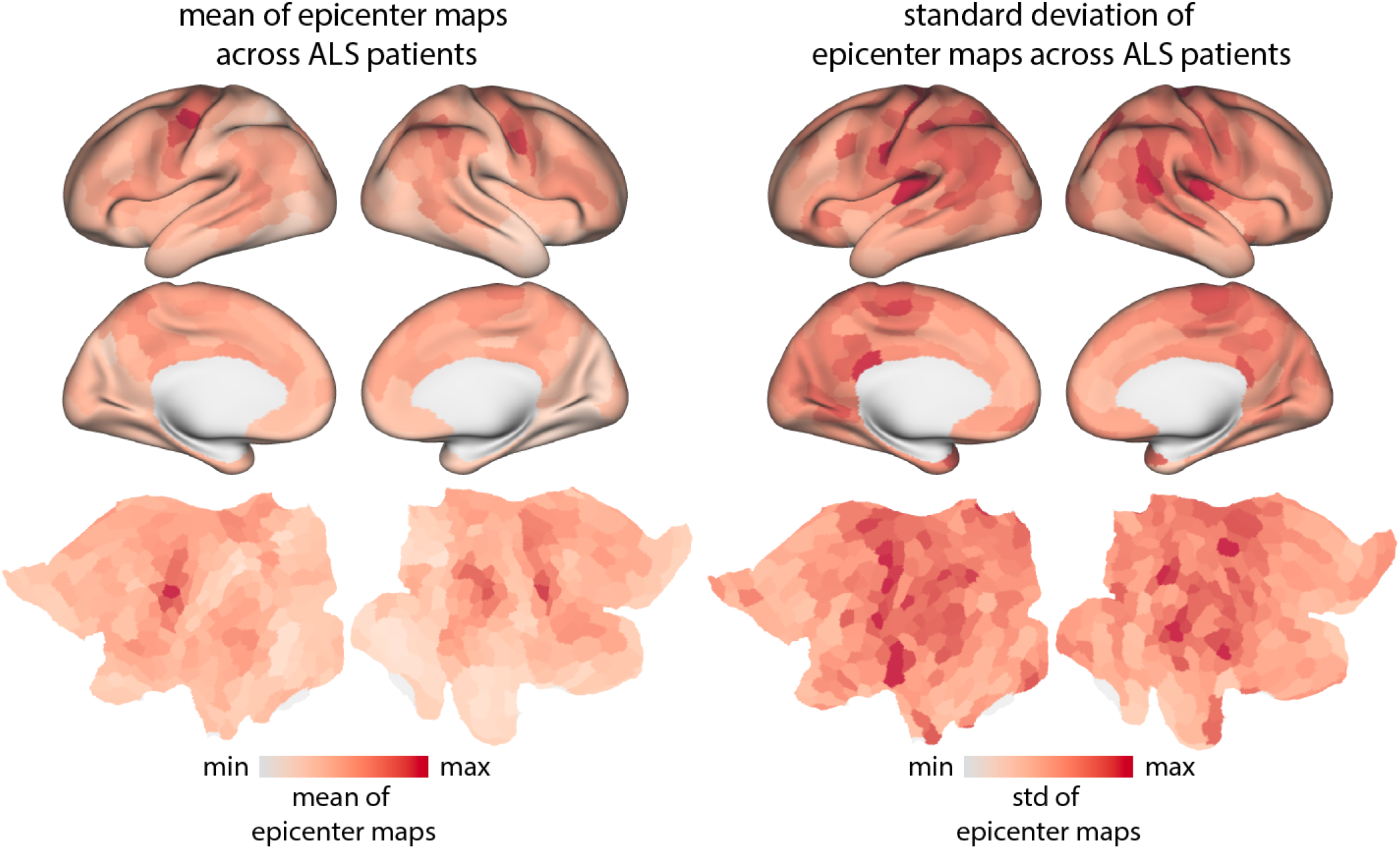
Map of mean and standard deviation of epicenter maps across individuals with ALS. Here the *w*-score map (after multiplying by −1) is used as a participant-specific atrophy map. Each cortical node is then assigned an epicenter likelihood value using the atrophy ranking method (see *Methods*). After estimating the epicenter likelihood map per participant, the map is normalized so that the highest-probability epicenter parcel scores 1, and the lowest scores 0. We average these normalized maps across all ALS participants to produce a mean epicenter map. The standard deviation for each parcel is also calculated based on these normalized values. The mean and standard deviation maps are shown on the fs-LR inflated and flat cortical surfaces.

**Figure S3.**
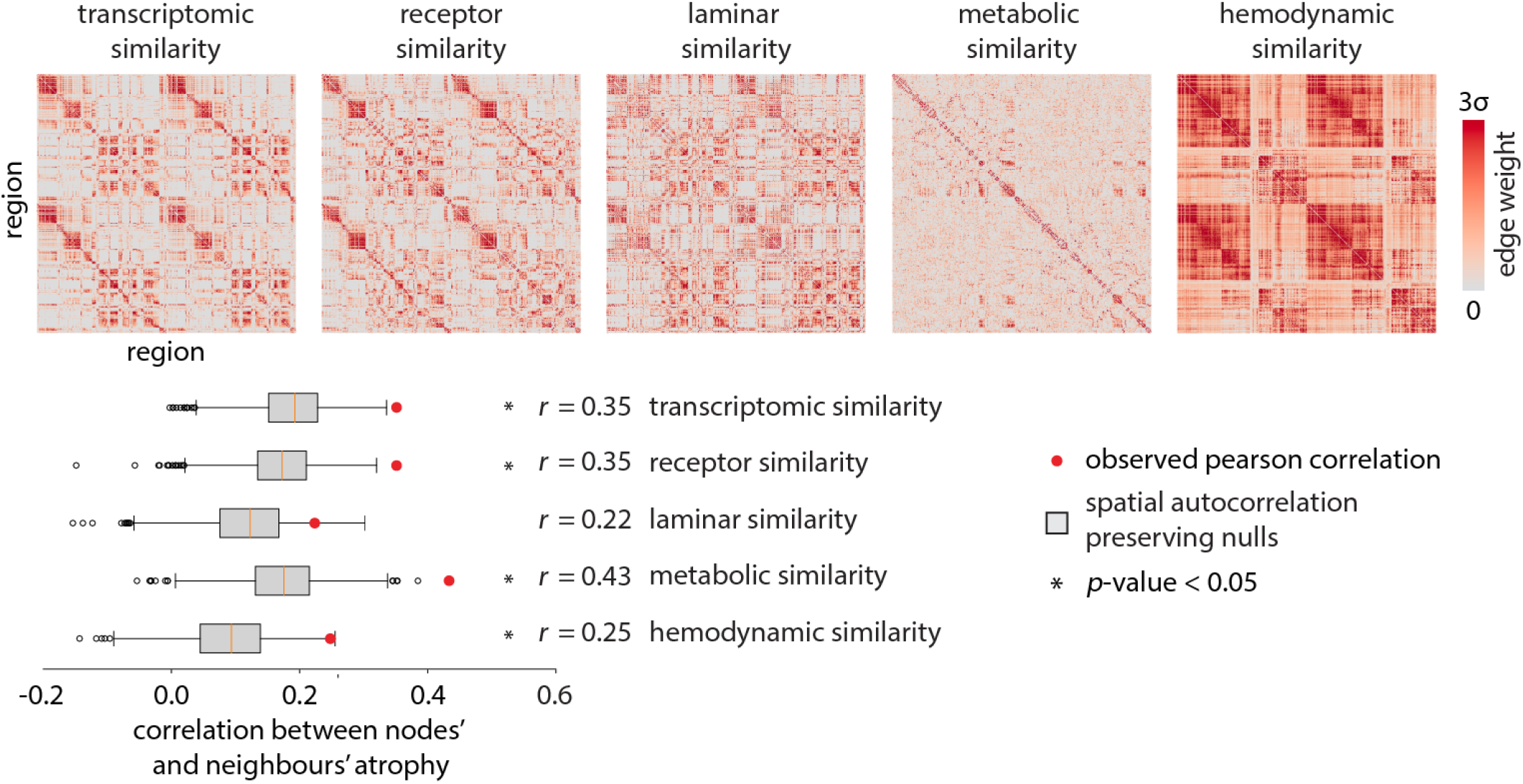
Inter-regional biological similarity shapes atrophy. The heatmaps show each biologically defined network according to the Schaefer-400 parcellation [111]. Negative-valued elements are excluded from all analyses. The networks are then used to calculate the node-neighbour atrophy correlations. The significance of node-neighbour correlation values is assessed with respect to spatial autocorrelation preserving spin tests. All networks except laminar similarity lead to significant node-neighbour correlation values (10, 000 repetitions; FDR corrected; gene expression similarity: *p*_spin_ = 1.33 × 10^−2^, receptor similarity: *p*_spin_ = 1.25 × 10^−2^, metabolic similarity: *p*_spin_ = 4.99 × 10^−3^, hemodynamic similarity: *p*_spin_ = 2.75 × 10^−2^).

**Figure S4.**
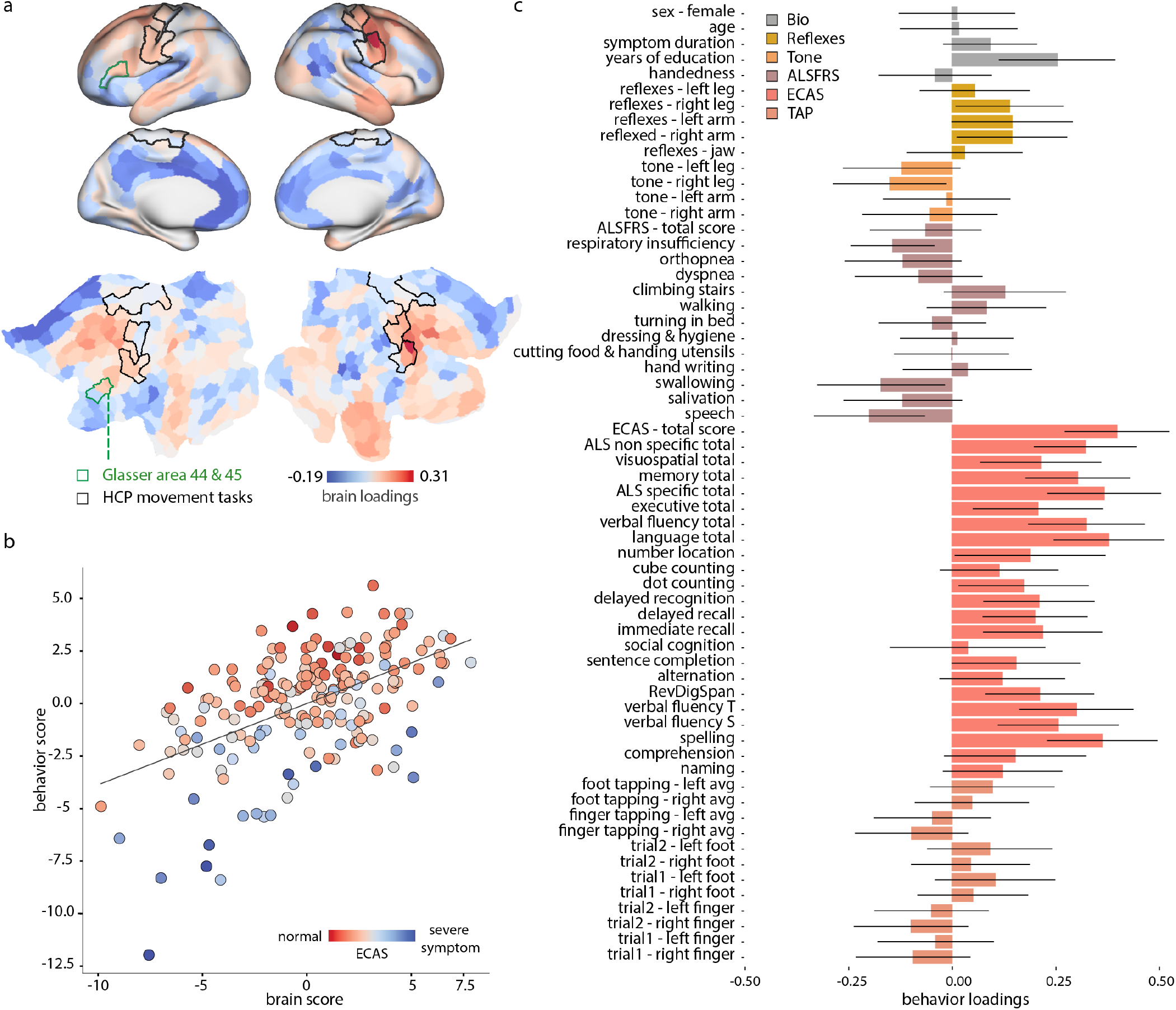
Second latent variable from a PLS analysis to relate individual epicenter maps with clinical and behavioral measures. (a) Brain loadings are shown on the fs-LR inflated and flat cortical surfaces. Regions demarcated by the black border are those with the greatest effect sizes (Cohen’s d effect size greater than 1) in the group-average activation map from S1200 Human Connectome Project package for the movement task contrasts [12]. Regions demarcated by green borders showcase areas 44 and 45 from the Glasser parcellation [51]. These regions, specifically in the left hemisphere [105], correspond the Broca’s area [16]. (b) The scatter plot visualizes the individual participants’ brain scores versus behavioral PLS scores (Pearson correlation coefficient, *r* = 0.52; Spearman correlation coefficient, *r* = 0.48); each participant’s score is colored based on the ECAS total score. (c) The bar plot visualizes the behavioral/clinical measures’ loadings. The contribution (effect size) of individual variables is assessed by bootstrap resampling (1, 000 repetitions).

**Figure S5.**
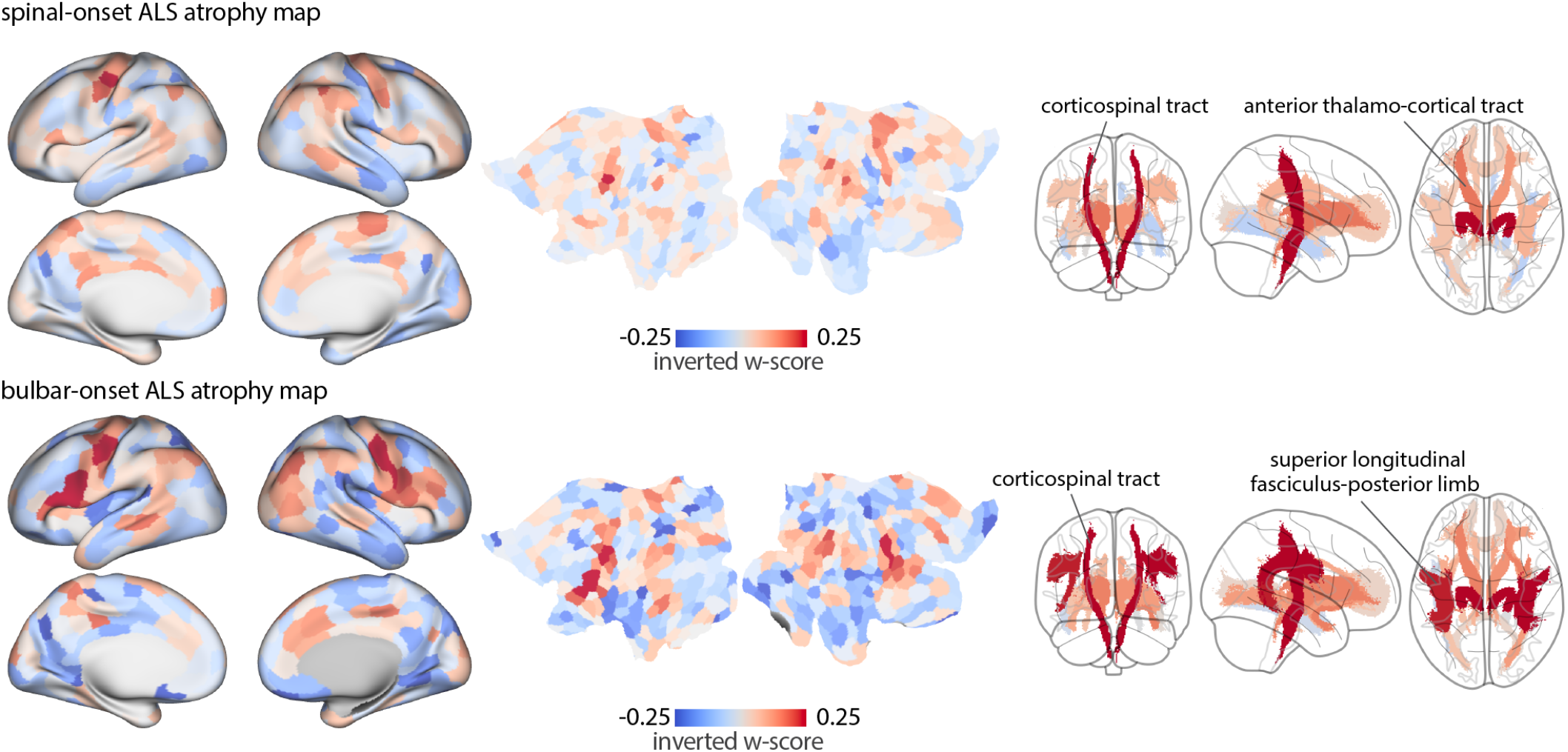
Map of ALS atrophy stratified by type of disease onset. Measuring the atrophy within atlas-based defined tracts [60, 146] showed that while corticospinal tract is involved in both spinal-(FDR corrected; left: *p* = 1.45 × 10^−9^, *t*-statistic= 7.50; right: *p* = 7.72 × 10^−10^, *t*-statistic= 7.74) and bulbar-onset ALS (FDR corrected; left: *p* = 1.72 × 10^−3^, *t*-statistic= 5.26; right: *p* = 5.52×10^−4^, *t*-statistic= 4.40). For patients with the spinal-onset of the disease, the anterior thalamic region is also significantly atrophied (FDR corrected; left: *p* = 4.31 × 10^−5^, *t*-statistic= 5.34; right: *p* = 9.62 × 10^−4^, *t*-statistic= 4.55). The involvement of anterior thalamic radiation in ALS has been reported in the cited references [37, 93]. In patients with the bulbar-onset of the disease, significant atrophy is also observed in the superior longitudinal fasciculus tract (FDR corrected; left: *p* = 1.72 × 10^−3^, *t*-statistic= 4.88; right: *p* = 5.34 × 10^−3^, *t*-statistic= 4.40). This tract is known to play role in speech [140] and language functions [95]. Decrease in fractional anisotropy of the superior longitudinal fasciculus tract in bulbar-onset ALS patients has been reported in previous studies [18, 127].

**TABLE S1.** Parameters of the agent-based SIR model. For more information on the model equations refer to [157].

| Notation | Name | Expression or value | Explanation |
| --- | --- | --- | --- |
| $\Delta t$ | time step | $\Delta t = 0.02$ | Time increments in the simulations |
| $\rho_i$ | the probability of remaining in region $i$ | $\rho_i = 0.99$ for all $i$ | Agents in region $i$ have equal probability of remaining in region $i$ or exiting region $i$ per unit time |
| $k_1$ | weight of atrophy accrual due to accumulation of misfolded agents | $k_1 = k_2 = 0.5$ | The contribution of native misfolded agents to total atrophy growth |
| $k_2$ | weight of atrophy accrual due to deafferentation | $k_1 = k_2 = 0.5$ | The contribution of deafferentation to total atrophy growth |
| $w_{ij}$ | Connection strength between regions $i$ and $j$ | Log-transform of streamline density | |
| $l_{ij}$ | Connection length between regions $i$ and $j$ | Euclidean length of streamline fibers | |

